# Scalable and semi-automated Boolean network inference from multi-condition single-cell transcriptomes with scBOLT

**DOI:** 10.64898/2026.10.02.756175

**Authors:** Théo Roncalli, Elisabeth Remy, Loïc Paulevé

## Abstract

Predictive logical models have made substantial contributions to biomedical research by facilitating the identification of regulatory mechanisms driving biological processes. Yet, their construction remains a major bottleneck, as it typically relies on extensive prior knowledge at the genome scale. The emergence of high-throughput omics technologies has opened new opportunities for data-driven logical model construction by providing high-resolution observations of biological states. In parallel, formal constraint-based approaches have emerged, reformulating executable model inference as a constraint-satisfaction problem. However, transforming omics observations into biologically meaningful Boolean state abstractions and dynamical constraints suitable for logical synthesis engines remains challenging, particularly for poorly understood biological systems and multi-condition experimental settings. To address these challenges, we introduce scBOLT, a semi-automated framework for Boolean constraint engineering and logical model inference from multi-condition single-cell transcriptomes. Beyond these methodological contributions, scBOLT substantially improves computational tractability, enabling the inference of Boolean networks comprising several hundred components within practical runtimes.

## 1 Introduction

Deciphering cellular processes requires identifying regulatory mechanisms driving cell fate decisions, with applications ranging from cellular differentiation and reprogramming to drug target discovery [31,96]. To investigate such mechanisms, a wide range of analytical formalisms have been developed. Among them, executable logical models have aroused increasing interest as their coarse-grained representation enables the scalable modelling of genome-wide regulatory systems while preserving key dynamical properties.

Boolean Networks (BNs) are widely employed executable models in systems biology, owing to their ability to represent regulatory interactions without requiring extensive kinetic parameterization [42,90]. They have been successfully applied to model a wide range of biological phenomena, including cell death decisions [97,11], tumorigenesis [24,78,64], and cell differentiation [59,67,1,20,70]. A BN is defined by a finite set of components, each governed by a logical rule and associated with a binary state, conventionally denoted 0 or 1. In transcriptome-derived scBOLT abstractions, these values represent off- and on-like transcriptomic states, respectively, rather than direct measurements of protein activity. The temporal evolution of states is determined by an update mode specifying how component states evolve according to these rules.

Such models are traditionally constructed using forward, literature-based modelling approaches that rely on expert curation and trial-and-error refinement [11,20,82]. Over the past decade, high-throughput sequencing technologies have reframed biological modelling as an inverse problem, where regulatory models are inferred directly from omics data [85,64]. However, despite the development of an ecosystem of tools for extracting biological information from omics datasets, reconstructing executable regulatory models remains challenging due to the combinatorial complexity of genome-scale regulatory networks.

Early approaches have nevertheless provided partial answers to this challenge. In particular, considerable efforts have been devoted to the inference of relationships between biomolecular entities through the analysis of statistical dependencies between expression profiles. A major difficulty, however, lies in determining whether statistical associations reflect direct regulatory relationships or instead arise from regulatory cascades and shared upstream regulators. ARACNE [56] and CLR [25] were among the first methods to exploit mutual-information-based measures for disentangling direct from indirect associations in large microarray expression compendia. Yet, such associations do not provide information about the directionality of the inferred relationships. Consequently, alternative microarray-based approaches emerged to identify candidate regulator-target relationships. GENIE3 [40] notably framed interaction reconstruction as a collection of independent regression problems whose predictions were aggregated through tree-based ensemble methods, enabling the inference of directed regulatory dependencies. While previous methods remained largely restricted to static descriptions of molecular systems, subsequent methods such as Inferelator [10] and dynGENIE3 [39] further extended these modelling approaches by incorporating temporal information to infer dynamical regulatory models from time-series measurements. The advent of next-generation sequencing (NGS) technologies substantially expanded the scope of regulatory network inference by enabling the characterization of increasingly complex and heterogeneous cellular populations. Building upon these advances, SCENIC [2] combined co-expression inference with cis-regulatory motif analysis to identify context-specific regulatory interactions from single-cell transcriptomic data. Collectively, these approaches have substantially advanced the data-driven inference of gene regulatory networks (GRNs). Yet, inferred GRNs primarily describe relationships between biomolecular entities rather than the logical rules governing how multiple signals are integrated to generate cellular behaviours.

To address this limitation, several approaches have sought to infer BNs directly from omics data. Many rely on heuristic optimisation strategies to cope with the combinatorial complexity of logic rule inference, often in the context of specific clinical applications. For instance, DrugLogics [26] infers executable logical models constrained by a prior GRN and upstream knowledge of drug-gene interactions, using a genetic algorithm that maximises agreement between simulated steady states and data-derived stable states, enabling drug synergy prediction and patient-specific modelling. In contrast, BONITA-RD [72] infers BNs by applying a genetic algorithm based on network propagation with fuzzy logic values. Because genetic algorithms may struggle to find globally optimal solutions when the prior influence graph is dense, BONITA-RD couples it with a node-wise local search. This strategy optimises logic rules for each gene while keeping remaining rules unchanged. Nevertheless, the method remains computationally intensive, restricting the search space to at most three transcription factors per gene.

As an alternative, IQCELL [36] combines statistical network reconstruction with formal constraint solving to infer BNs without requiring a previous definition of the possible interactions between macromolecules. First, it infers a GRN using gene mutual information, then ranks genes along a pseudotime trajectory to identify causal interactions. Logical rules are then constructed using a satisfiability modulo theory engine (Z3) [21]. However, this approach introduces asymmetric linear influences where top-ranked genes are not influenced and lower-ranked genes exert limited influences, preventing the learning of some network motifs. Additionally, the inference procedure relies on a single trajectory, restraining its applicability to a limited subset of biological processes.

More recently, formal synthesis has emerged as an alternative to numerical optimisation for inferring executable models. Rather than relying on objective-driven searches to identify BNs consistent with experimental observations, these methods formulate BN inference as a combinatorial constraint-solving problem, enabling the exhaustive enumeration of BNs satisfying a set of formal constraints. Boolean Network Sketches [6] proposes a symbolic method for enumerating exact solutions with respect to a set of formal restrictions. The constraints encompass partially specified Boolean functions, monotonicity properties, mandatory regulators, and dynamical properties expressed in a hybrid extension of computation tree logic (HCTL). However, specifying network properties through temporal logic constraints may remain difficult. Alternatively, BoNesis [17,18,16] provides a domain-specific language for expressing dynamical constraints without directly manipulating temporal logic formulae. Declarative specifications are translated into an Answer-Set Programming (ASP) logic problem, enabling the synthesis of ensembles of BNs satisfying dynamical Boolean properties specified over partially defined states, within a predefined domain (e.g. a GRN, a set of BNs, or a partially defined BN). The framework supports a broad spectrum of dynamical properties related to trajectories, attractors and mutations, such as the existence or exclusion of transitions between cellular states. To satisfy these properties, BoNesis relies on the Most Permissive (MP) update mode, capturing all behaviours achievable by any underlying quantitative model consistent with the same logical structure [13,74].

Unlike numerical optimisation-based inference methods, which can trade off between competing objectives, formal synthesis approaches require all specified constraints to be simultaneously satisfiable. As a result, the combination of extensive candidate domains and numerous biological constraints may lead to unsatisfiable problems, i.e., having no solution. Recent developments in BoNesis provide combinatorial optimisation procedures for identifying maximum-cardinality jointly satisfiable component sets, thereby facilitating the construction of large candidate domains. In addition, BoNesis can maximise the cardinality of sets of *strong constants*, i.e., component sets whose members, within a single compatible realisation, jointly admit constant Boolean functions matching their respective values in every completed configuration. Removing one such optimal set maximally reduces the regulatory domain under the active constraints by fixing and eliminating the components in that selected strong-constant set, enabling the enumeration of sparsest compatible BNs as defined by the strong-constant criterion.

While constraint-based frameworks provide powerful tools for BN synthesis, translating omics data into biologically relevant Boolean constraints remains largely unresolved. To address this challenge, a first workflow was proposed by Herault et al. [35], combining single-cell transcriptomic data with BoNesis-based logical synthesis to model early hematopoiesis aging. While largely data-driven, the workflow still relied on substantial expert intervention, notably for regulatory component selection and the characterisation of biologically relevant Boolean cellular states, resulting in a compact model restricted to 15 regulatory components selected from an initial network of 133 transcription factors.

More recently, Chevalier et al. [15] proposed a methodology (Fig. 1) explicitly dedicated to constructing the inputs required by a logical synthesis engine from single-cell and bulk transcriptomic data. Applied to hematopoietic differentiation, the resulting inputs enabled the use of BoNesis to infer ensembles of BNs reproducing expected cellular differentiation processes. First, a prior GRN assembled from curated regulatory databases [28] was used as the domain restraining the search space of candidate BNs, specifying which activators and inhibitors are allowed. At single-cell resolution, an elastic principal graph was inferred in a spectral embedding [5] using STREAM [14], and cells associated with its nodes were used to define macrostates representing groups of cells sharing similar phenotypic profiles. For each macrostate, a partially defined Boolean state was estimated with PROFILE [4], where only the most discriminative features were assigned a Boolean value. Finally, trajectory relationships between macrostates were derived from the inferred lineage structure and prior biological knowledge, and used to formulate dynamical constraints describing admissible transitions between cellular states. Although successfully reproducing expected biological behaviours, the proposed workflow for single-cell data was demonstrated on hematopoiesis, a well-established differentiation system exhibiting a largely consensual lineage structure. As a result, both macrostate construction and trajectory inference benefit from substantial biological knowledge, making the generalisation of the workflow to more complex systems less straightforward.

**Fig. 1:**
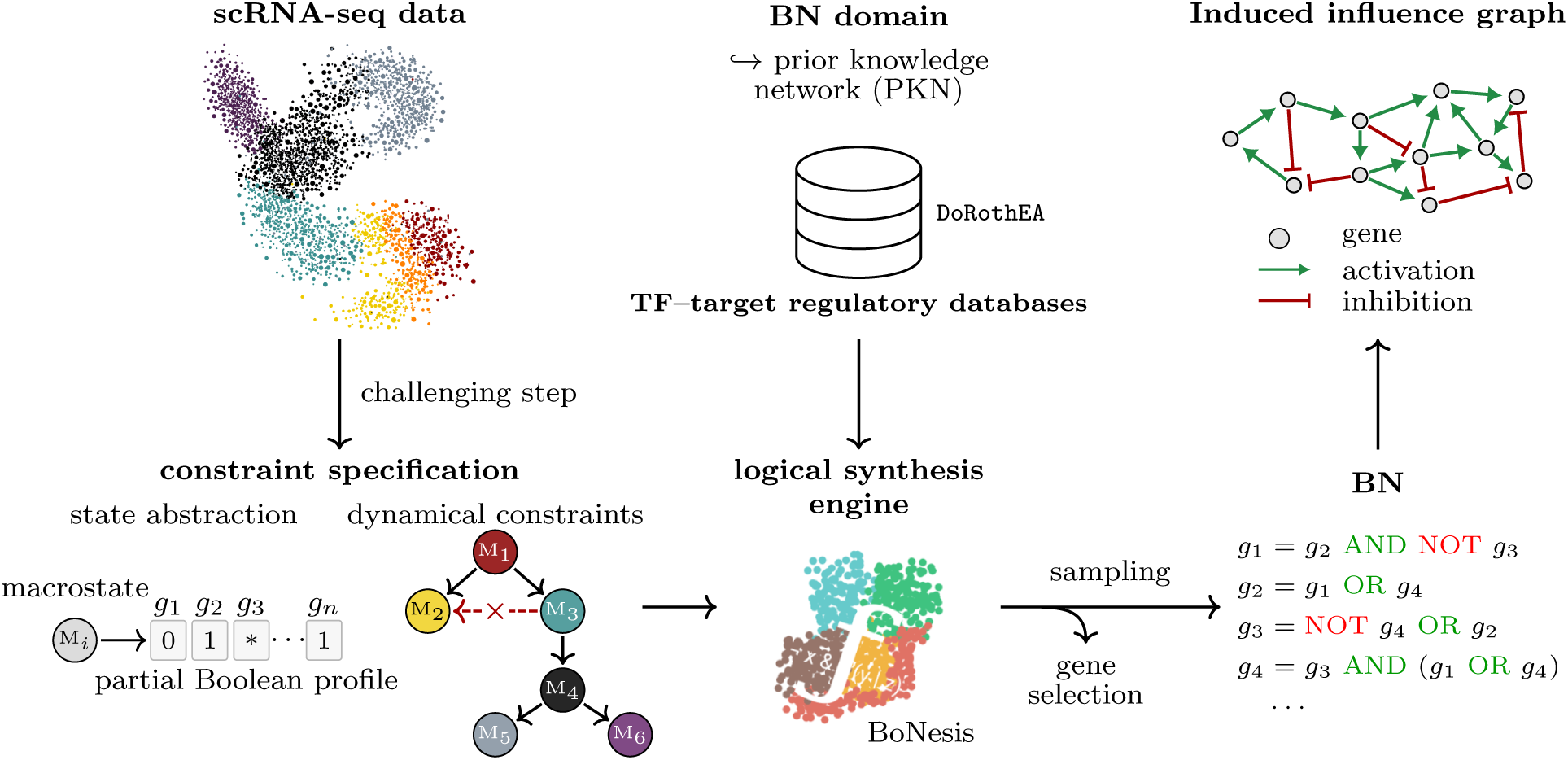
General methodology for single-cell Boolean modelling. Single-cell transcriptomes are summarised into macrostates (M*i*) and partially defined Boolean profiles, where 0, 1 and *∗* denote low, high and undefined transcriptomic states, respectively. Black and crossed red arrows indicate admissible and excluded transitions between macrostates. In parallel, prior TF–target regulatory databases define the regulatory domain and its signed influence graph. These inputs are combined as Boolean constraints for BoNesis-based logical synthesis.

As deriving Boolean constraints from omics observations remains challenging, we developed scBOLT (single-cell BOolean network Learning from multi-condition Transcriptomes), a framework built upon BoNesis to infer ensembles of sparsest BNs from scRNA-seq data. First, scBOLT facilitates transcriptome-driven Boolean constraint engineering by providing an abstraction layer between single-cell observations and exact logical synthesis engines, reducing the amount of expert intervention required throughout the modelling process. Second, scBOLT facilitates the modelling of poorly documented and non-canonical biological systems, reducing the reliance on extensive prior knowledge and improving the generalisability of Boolean constraint engineering workflows. Third, scBOLT supports multi-condition modelling, enabling the joint inference of BNs from multiple experimental settings while mitigating condition-specific and individual-specific biases. Once Boolean constraints have been formulated, exact BN inference may become intractable for highly constrained genome-scale synthesis problems due to their combinatorial complexity. Fourth, scBOLT substantially improves computational scalability through staged and adaptive decomposition of the underlying ASP optimisation problems, enabling constraint-driven identification of large and informative candidate domains. Together, these contributions extend constraint-based logical model inference to a broader range of biological systems and experimental settings, while expanding its computational reach to BNs comprising several hundred components within practical runtimes.

Beyond these methodological contributions, scBOLT is delivered as a modular software framework. It provides multiple analytical strategies, decision-support analyses and interoperable entry points, allowing users to integrate external analyses and adapt the framework to diverse modelling scenarios. Rather than enforcing a fully automated pipeline, a few stages are deliberately left configurable, acknowledging that modelling choices ultimately depend on the biological questions being addressed. Reproducibility is supported through configuration-driven execution, controlled dependency management, versioned external resources and controlled computational environments. Together, these mechanisms support reproducible single-cell analyses within controlled Linux x86-64 environments. Within the reference Intel Meteor Lake environment, strict numerical identity was verified across repeated runs for all tested single-cell analysis stages. Cross-architecture identity was additionally validated for a subset of the resulting outputs across qualified Intel Meteor Lake and AMD Zen 3 environments using matching SHA-256 digests; the exact validation scope is detailed in Supplementary Note 3. BN inference provides a distinct guarantee: solver certification establishes lexicographic optimality of the configured objective vector, although tied optimal component sets may exist. For larger inference problems, scBOLT provides time-limited adaptive optimisation strategies that improve tractability but may return hardware-and schedule-dependent incumbents, enabling the inference of BNs comprising several hundred components.

We demonstrate complementary strengths of scBOLT through two case studies. The first investigates acute promyelocytic leukemia (APL) driven by the PLZF-RAR*α* fusion, a rare and therapy-resistant variant of the disease [50,34,22]. Unlike classical PML-RAR*α* APL, this PLZF-RAR*α* subtype remains poorly characterised despite decades of investigation and exhibits resistance to therapies that transformed the clinical management of APL, making it a particularly challenging biomedical setting for BN inference. The second revisits early hematopoietic differentiation, a well-established system previously modelled with BoNesis [15], providing a direct benchmark against the original monolithic gene-selection procedure and allowing scBOLT’s gains in computational tractability to be assessed.

## 2 Results

### 2.1 Workflow architecture

scBOLT relies on a semi-automated workflow to generate an ensemble of BNs reproducing cellular dynamics from multi-condition single-cell transcriptomic data. It uses BoNesis as its core inference framework, where BN synthesis is driven by structural constraints restricting the model search space and dynamical Boolean constraints. A central challenge lies in translating single-cell transcriptomic data into meaningful dynamical constraints. scBOLT addresses this by identifying macrostates corresponding to phenotypic cell populations, which are subsequently binarised into partially defined Boolean states to restrain structural properties. Relationships between macrostates, deduced from biological interpretation and trajectory analyses, are encoded as dynamical constraints, enabling behaviours consistent with observed dynamics.

Figure 2 provides an overview of the workflow, summarising the main steps from data preprocessing to BN inference. Most steps are fully automated, but some, highlighted in red, require user intervention. In addition to the conceptual workflow, scBOLT exposes its analytical and modelling modules through a unified command-line interface, allowing users to execute individual stages, inspect and reuse intermediate outputs, and reproduce analyses from recorded configurations. From input datasets in FASTQ format, it generates output models in the standard .bnet format. A detailed diagram of the pipeline, highlighting each module and its role across the workflow, is provided in Figure 7. The underlying methods are described in Section 4.

**Fig. 2:**
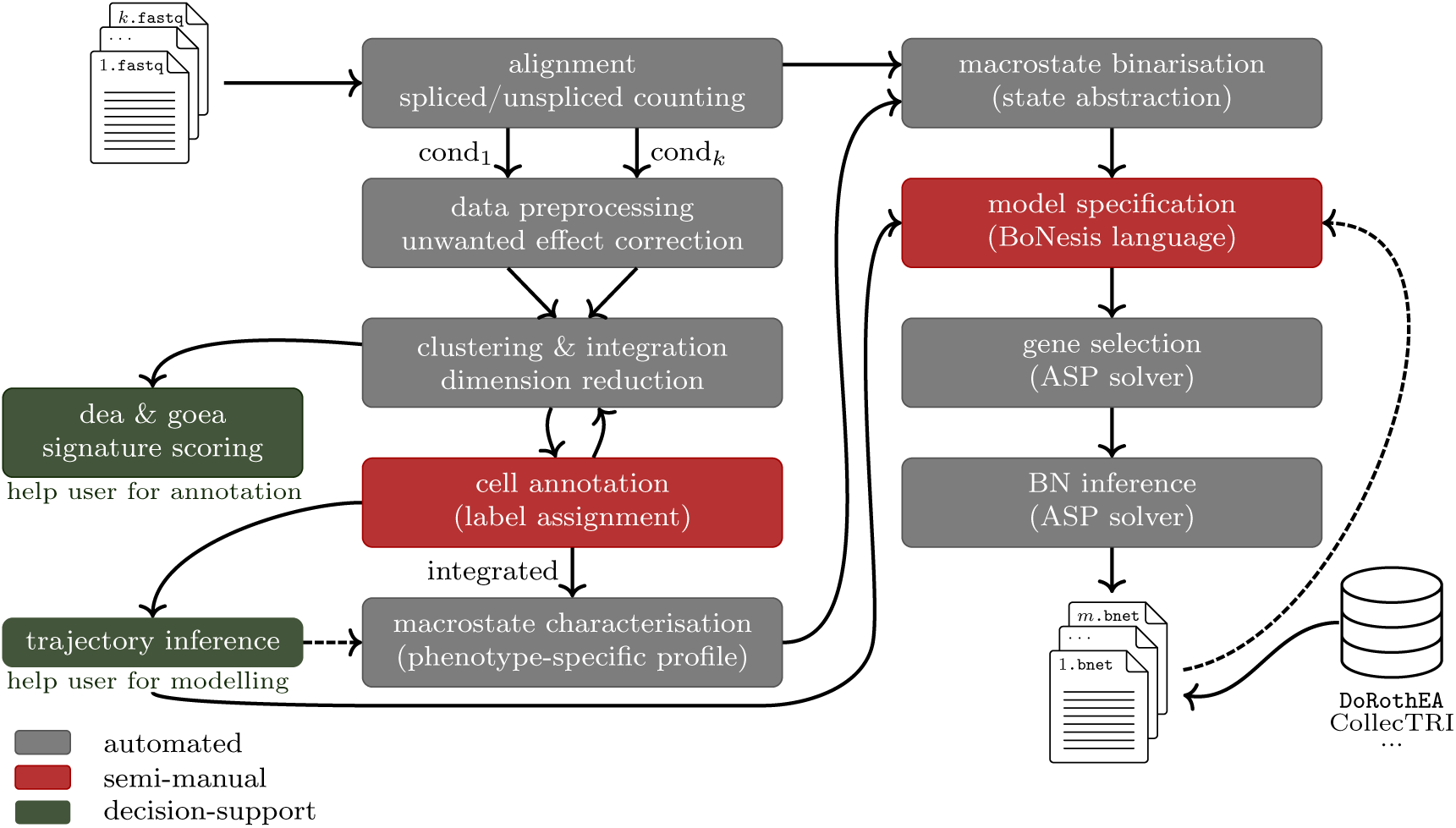
scBOLT workflow. Main steps from scRNA-seq input data (FASTQ format) to output BNs (.bnet format). Computational steps are either automated (grey) or require user input (red), with optional decision-support analyses (green). Note that the macrostate binarisation step requires particular care, as multiple strategies are available and the default choice may not always be optimal.

The workflow begins with standard preprocessing and integration steps applied to single-cell transcriptomic data, preparing them for downstream operations. This includes raw data processing (read alignment to a reference genome and spliced/unspliced quantification), followed by dataset-specific preprocessing steps encompassing normalisation, filtering, and optionally the removal of cell cycle phase variation effects. These steps aim to capture biologically relevant signals rather than technical or confounding variability. Clustering, integration and dimensionality reduction techniques are then performed jointly across datasets to identify cellular populations with similar gene expression profiles across conditions, while reducing data complexity. In particular, dataset integration aligns shared phenotype-related populations across wet-lab experiments, mitigating batch effects and preventing condition-specific biases from propagating to downstream modelling.

scBOLT then supports cell annotation at the cluster level, which help define which biological processes will be modelled and to guide the subsequent specification of dynamical constraints, as identified phenotypes may already suggest plausible state transitions. This step requires user intervention, as automated annotation methods remain unreliable in poorly characterised biological contexts and may fail to capture relevant phenotypic distinctions. To support this process, scBOLT provides decision-support tools, including differential expression analyses (DEA), gene ontology enrichment analyses (GOEA) [43], and signature scoring [94,12].

In more complex settings, however, transcriptomic dynamics cannot be deduced from annotation alone, limiting its applicability to well-documented processes. scBOLT therefore provides trajectory inference tools to characterise temporal relationships between cell states for guiding the formulation of dynamical constraints, including RNA velocity [8] and CytoTRACE 2 potency-based analyses [30,41]. RNA velocity estimates a vector field over the phenotypic manifold from splicing kinetics, while potency quantifies differentiation potential from gene expression levels. However, velocity-based methods rely on simplified kinetic models and require sufficient curvature in the spliced–unspliced phase space to distinguish up- from down-regulation [9]. For example, transcriptional boosts observed in erythroid differentiation can yield reversed velocity estimates, where increasing gene expression is misinterpreted as downregulation [75,84,3]. Additionally, incomplete sampling of dynamic processes can lead to ambiguous velocity directions [9].

To complement the specification of dynamical constraints, scBOLT constrains state properties using a two-step procedure: (i) defining macrostates as groups of cells sharing the same phenotypic patterns for each experimental condition, and (ii) computing a partially binarised vector for each macrostate, indicating low and high transcriptomic states. scBOLT provides several approaches for computing macrostates, including CellRank [45], STREAM [14] and COTAN [27]; the appropriate choice depends on the biological context. CellRank identifies macrostates from velocity, potency and transcriptional similarity relationships between cells. Although used by default, its performance may degrade when velocity assumptions are not satisfied. COTAN derives macrostates from pairwise gene co-expression patterns inferred from zero UMI counts, making it particularly suited for highly sparse datasets. However, as a method designed originally for clustering, it may generate many partitions, requiring a deep understanding of the dataset to identify biologically meaningful macrostates. STREAM, originally designed for trajectory inference, derives macrostates from branching structures inferred in low-dimensional embedding spaces. Its performance may, however, degrade when the embedding space exhibits low topological complexity, in particular when cells form a homogeneous cloud without clear branching.

To address these limitations, we propose an additional macrostate identification strategy: *k*-nearest-neighbour-based subclustering (KNNSC). It refines precomputed clusters into fine-grained phenotypic populations by exploiting local relationships in reduced embedding spaces, thereby remaining robust to distortions of global nonlinear distances. It can prioritise either central or peripheral cellular populations within clusters. However, both strategies may become less informative when a single cluster contains spatially separated cells, since the cluster’s centroid may no longer accurately represent local cellular organisation. Although KNNSC ensures graph connectivity across clusters, it does not explicitly resolve such intra-cluster multimodal structures.

Following macrostate identification, scBOLT performs binarisation to derive partially defined Boolean vectors at the macrostate level by aggregating single-cell gene expression counts. This step assigns genes a low (0), high (1), or undefined transcriptomic state within each macrostate, thereby directly constraining the inferred BNs. These coarse-grained assignments reflect relative RNA abundance and do not directly measure protein activity. As no consensus exists in the field, scBOLT provides diverse binarisation strategies. A distribution-based approach proceeds in two steps: (i) assigning partially defined Boolean values at the single-cell level using scBoolSeq [55], which infers gene expression distributions and applies distribution-specific binarisation rules, and (ii) aggregating them at the macrostate level using a voting scheme. Nevertheless, scBoolSeq is limited in sparse datasets with high dropout rates, as it cannot distinguish true biological absence of expression from technical dropout at the single-cell level. To overcome this limitation, a DEA-based approach directly binarises macrostates, averaging out technical dropout effects across cells. It identifies significantly over- or under-expressed genes using differential-expression tests, with the Wilcoxon rank-sum test as default and two variants of Welch’s *t*-test also supported, leaving non-significant genes undefined. Finally, both approaches can be combined within a consensus framework to improve robustness and ensure consistency across methods.

Using the partially Boolean-encoded macrostates, scBOLT then proceeds to the specification of dynamical constraints using the domain-specific declarative language BoNesis [16], a key step that remains manual to allow users to tailor the model to their specific objectives. Dynamical constraints are decomposed as follows. Semantics-invariant constraints are derived from macrostates, including the definition of configurations (fully binarised states matching macrostate profiles), static constraints between configurations (e.g. equality or inequality), and attractor properties such as fixed points or trap spaces. Trajectory constraints are guided by annotation and trajectory analyses, and encode temporal relationships between states, including (non-)reachability between configurations and reachable attractors.

Then, the inference task consists in identifying an ensemble of BNs that satisfy user-defined Boolean constraints while being restricted to a predefined regulatory domain. With scBOLT, this domain corresponds to a database-derived GRN [28,63,65] defining the space of candidate activating and inhibitory interactions. Nevertheless, inferring BNs suffers from a combinatorial blow-up, as the number of admissible logical rules for a gene increases exponentially with the number of its regulators, and the cost of ASP-based verification of a BN also grows with the number of its components. To make inference tractable at genome scale, scBOLT first maximises a jointly satisfiable component set under the imposed dynamical constraints, then removes an optimised set of strong constants from the propagated domain using optimisation features provided by BoNesis. Components omitted by the joint optimisation are not thereby shown to be individually incompatible. Subset-minimal BNs are then inferred using combinatorial solving strategies offered by BoNesis, enabling the identification of regulatory mechanisms driving the observed cellular dynamics.

### 2.2 Case study: treatment-resistant acute promyelocytic leukemia

Acute Promyelocytic Leukemia (APL) is an acute myeloid leukemia characterised by a block in early granulocytic differentiation, leading to the accumulation of immature promyelocytes in the bone marrow. It is driven by de novo chromosomal translocations involving the retinoic acid receptor alpha (RAR*α*) locus. These alterations impair transduction of the retinoid signalling pathway, promoting transcriptional repression of genes required for granulocytic maturation [22]. APL is most commonly associated with the t(15;17)(q24;q21) translocation, which generates the PML-RAR*α* fusion oncoprotein and accounts for approximately 98% of APL cases [29]. The introduction of all-trans retinoic acid (ATRA) [38] and arsenic trioxide (ATO) [86] has transformed the clinical management of PML-RAR*α* APL [52]. ATRA and ATO respectively target the RAR*α* and PML moieties of the PML-RAR*α* fusion oncoprotein, promoting its ubiquitin–proteasome-mediated degradation and restoring granulocytic differentiation [22]. These therapies have converted a previously fatal disease into one of the most successful examples of targeted cancer therapy [22,29]. However, the most common non-PML-RAR*α* variant, arising from the t(11;17)(q23;q21) translocation that generates the PLZF-RAR*α* fusion oncoprotein, is refractory to ATRA- and ATO-based therapies [29]. Unlike PML-RAR*α* APL, ATRA-induced degradation of the PLZF-RAR*α* fusion oncoprotein fails to clear self-renewing leukemic cells, resulting in persistent disease despite partial granulocytic differentiation [34,77]. Consequently, PLZF-RAR*α* APL remains poorly characterised after decades of investigation, making it a particularly challenging benchmark for data-driven BN inference.

To investigate the regulatory mechanisms underlying therapeutic resistance in PLZF-RAR*α* APL, we analysed a multi-condition scRNA-seq dataset generated at the CRCM laboratory (Marseille, France) [77], following the experimental protocol of He et al. [33]. Cells were isolated from the bone marrow of mice transplanted with PLZF-RAR*α* leukemic cells and either left untreated or treated in vivo with all-trans retinoic acid (ATRA) for seven consecutive days. The scBOLT framework was applied to infer ensembles of BNs from these single-cell transcriptomic data. While most stages of the framework are fully automated, cell annotation, macrostate definition and constraint formulation rely on expert supervision, assisted by dedicated visualisation and decision-support tools.

#### Data description and preprocessing

Using the intermediate entry point supporting processed count matrices, publicly available multi-condition single-cell scRNA-seq data comprising untreated control and in vivo ATRA-treated leukemic cells were retrieved from the study of Poplineau et al. [77]. Following quality control, 5,285 control cells and 6,855 ATRA-treated cells were retained. Unwanted sources of variation, such as cell-cycle effects, were removed to focus on phenotype-related transcriptional signals, after which both conditions were integrated using BBKNN [76]. Dataset characteristics and principal single-cell settings are summarised in Supplementary Tables 2 and 3; additional methodological details are provided in Supplementary Note 1. The integrated data are visualised using a UMAP embedding (Fig. 3a), revealing both shared and condition-specific cell populations. The resulting geometric structure indicates that ATRA treatment substantially reshapes the transcriptomic landscape despite its incomplete therapeutic response. In particular, a prominent control-specific region shows no overlap with the treated condition and may correspond to ATRA-sensitive cell populations, consistent with previous observations [77].

**Fig. 3:**
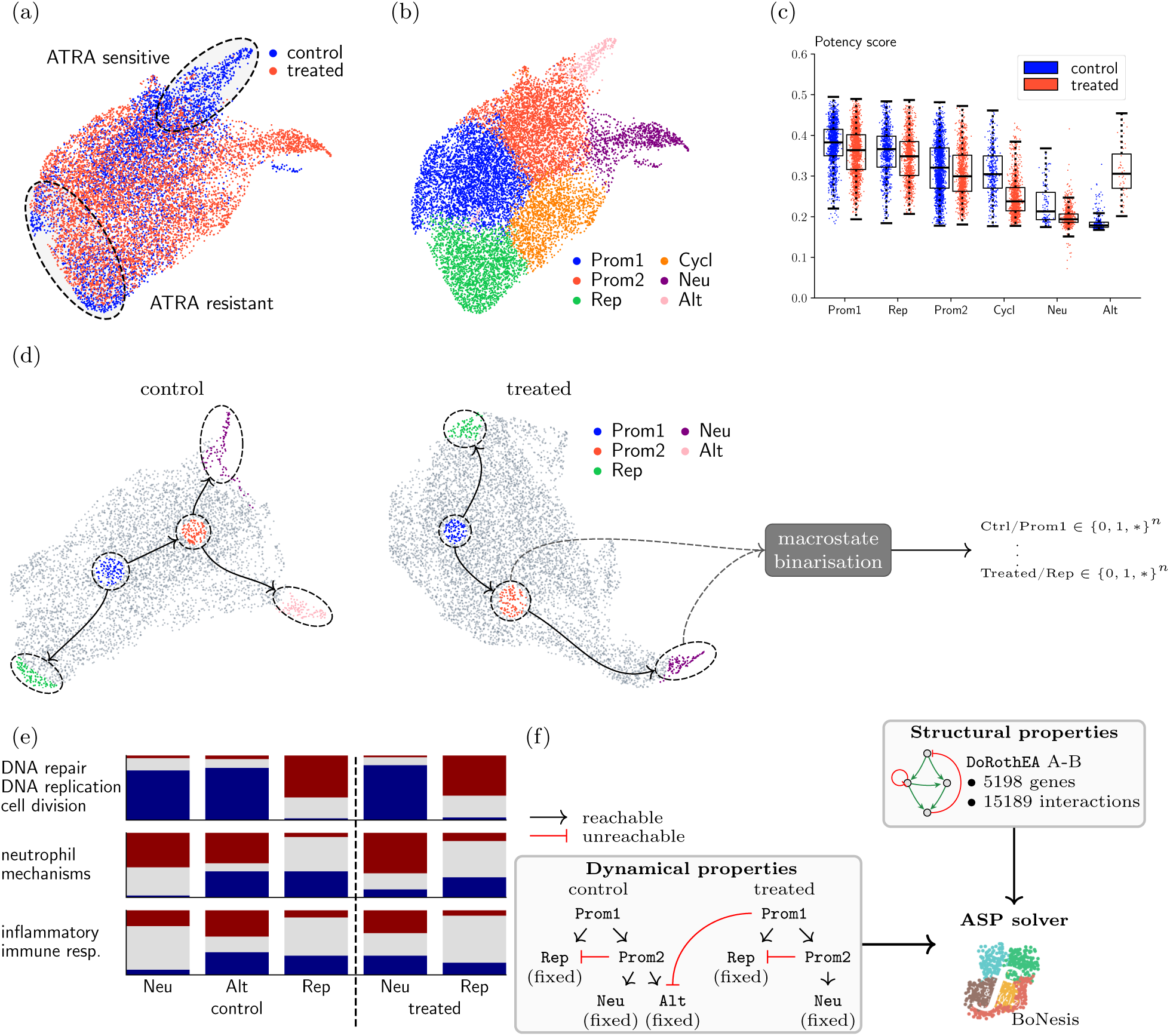
Single-cell transcriptome analyses of the PLZF-RAR*α* variant of APL. (a,b) 2D Uniform Manifold Approximation and Projection (UMAP) visualisation of integrated single-cell transcriptomes coloured by (a) condition and (b) manual cell annotation. Six clusters were identified: Prom1, Prom2, Rep, Cycl, Neu, and Alt. (c) Potency score distributions across clusters and conditions. Lower scores indicate more differentiated states. Condition-related groups are ordered by median developmental potential scores. (d) 2D UMAP embedding of the control and treated samples, showing cells coloured by macrostates identified with KNNSC, with arrows indicating the proposed relationships between macrostates. Macrostates are subsequently partially binarised into vectors in *{*0, 1*, ∗}^n^*. (e) Proportions of low (0), undefined, and high (1) transcriptomic assignments across terminal macrostates and conditions for selected biological process-associated gene sets, including proliferation-related processes (DNA repair, DNA replication, cell division), neutrophil functions (chemotaxis, activation, degranulation, and neutrophil-mediated immunity), and broader inflammatory or immune regulatory processes. Colours indicate low (blue), undefined (grey), and high (red) assignments. (f) Summary of the Boolean modelling framework, illustrating the input to BoNesis: the DoRothEA regulatory domain (structural restriction), together with semantics-invariant constraints (fixed points and trap spaces) and trajectory constraints (reachability and non-reachability under MP semantics).

#### Cell annotation

Cells were manually annotated (Fig. 3b) using differential-expression and enrichment analyses [43,94,12] performed on the integrated data (Supplementary Note 1), thereby enabling cross-condition annotation without per-dataset labelling. Six clusters were identified, consistent with previous findings [77]. The Neu and Alt clusters display granulocyte-associated features, whereas Rep and Cycl correspond to proliferative states, and Prom1 and Prom2 represent promyelocyte-like states lacking terminal myeloid programmes. While both Rep and Cycl share a common proliferative core, with enrichment in DNA replication, DNA repair and genome maintenance programmes, they exhibit distinct transcriptional identities. Rep is further distinguished by strong enrichment in RNA processing, spliceosomal assembly, ribosome biogenesis, translational processes and telomere maintenance, reflecting a biosynthetically active and dysregulated proliferative state. In contrast, Cycl is dominated by mitotic processes, including chromosome condensation, spindle organisation, kinetochore assembly, chromosome segregation and cytokinesis, consistent with a more canonical cycling programme. Notably, the Cycl cluster has a markedly higher relative abundance in the ATRA-treated condition than in control (18.0% versus 7.7%). In contrast, Rep remains present at comparable proportions across conditions, representing 17.5% and 18.4% of control and ATRA-treated cells, respectively (Supplementary Table 4). This persistence suggests that Rep may contribute to disease maintenance.

Both Neu and Alt clusters display immune- and inflammatory-related features, as well as migration-related processes potentially associated with egress from the bone marrow, and are enriched in quiescence-associated signatures [94], consistent with non-proliferative states. However, their functional profiles markedly diverge. Neu is characterised by enrichment in neutrophil-specific processes, including neutrophil chemotaxis, degranulation, phagocytosis and neutrophil-mediated immunity, as well as respiratory burst and components of the NADPH oxidase complex, and by the expression of canonical neutrophil markers (e.g. S100a8, S100a9, Ly6g, Ngp), supporting a coherent neutrophil-like programme. In contrast, Alt displays a broader immune and inflammatory programme associated with migration-, adhesion- and signalling-related processes, expresses eosinophil-associated genes (e.g. Epx, Prg3, Ear6, Alox15), and also expresses markers associated with endoplasmic-reticulum stress. This population likely reflects an altered, non-canonical differentiated lineage. It accounts for 250 of 5,285 control cells (4.7%) but only 52 of 6,855 ATRA-treated cells (0.8%), suggesting that ATRA-based therapy rewires differentiation trajectories, leading to the loss of the corresponding programme.

#### Trajectory inference

To guide specification of dynamical constraints, we investigated cell-state hierarchies through trajectory inference. This optional analysis is particularly useful here, as PLZF-RAR*α* APL-related biological processes remain poorly defined in the literature. Nevertheless, the assumptions underlying velocity-based methods are challenged in highly proliferative contexts. In particular, rapid expansion of promyelocytes induces heterogeneous transcriptional regimes, leading to inconsistent splicing-degradation dynamics that distort velocity estimates [44,8].

To overcome this limitation, we therefore relied on potency scores (Fig. 3c), which provide a robust, condition-independent measure of differentiation potential. Potency analyses indicate that cells are predominantly unipotent while still exhibiting a hierarchy. As expected, granulocytes (Neu, Alt) occupy the most differentiated states, although not fully mature as they remain in the bone marrow. A small subset of treated cells falls within the Alt cluster, likely reflecting clustering inaccuracies, as these cells localise near the Prom1 region in the embedding (Fig. 3b). Prom1 cells occupy the least differentiated states, potentially preceding other populations in cell-fate trajectories, and Prom2 cells also retain a precursor-like state but lie at a more advanced stage within the promyelocytic compartment, corresponding to an intermediate position along the differentiation hierarchy. While both Rep and Cycl are proliferative, Rep cells exhibit differentiation levels close to immature promyelocytes (Prom1), consistent with an early differentiation arrest characteristic of the leukemic process. Cycl cells display slightly higher differentiation levels, reflecting a more advanced cellcycle progression state. These relationships provide a basis for defining subsequent trajectory constraints, with cluster Prom1 as an initial state, and clusters Rep, Alt, and Neu as candidate terminal states.

#### Macrostate characterisation

Macrostates were then identified to obtain a reduced and interpretable representation of the cellular landscape, capturing key phenotypic patterns. This step proved particularly challenging in the APL dataset, as several macrostate inference strategies failed to provide suitable phenotypic abstractions. CellRank [45] was not retained, as its velocity-based kernel becomes unreliable owing to the strong transcriptional regime shifts that violate RNA-velocity assumptions. COTAN over-partitioned the data into numerous small populations, making biological interpretation difficult in an already complex cellular landscape. Likewise, STREAM was ill-suited because the UMAP embedding displays limited global branching structure.

We therefore applied KNNSC to the UMAP embedding, as its local topology-based strategy remains informative when the embedding is compact and exhibits limited global branching. This enabled the refinement of each cluster into fine-grained macrostates (Fig. 3d). Terminal states (Rep, Alt, Neu) were defined using the periphery-based strategy to capture trajectory endpoints, selecting cells furthest from neighbouring clusters. Conversely, precursor-like populations (Prom1, Prom2) were identified using the complementary centroid-based strategy, selecting cells closest to their cluster centroids. The Prom1 macrostate represents a particular case, as initial states are generally defined using the periphery-based strategy, whereas Prom1 was deliberately identified using the centroid-based strategy. This choice reflects both the transient nature of promyelocytes during hematopoiesis and the restricted scope of our dataset, which lacks upstream progenitor cells.

#### Macrostate binarisation

Macrostates were subsequently partially binarised using a consensus strategy combining intrinsic expression patterns with differential-expression signals across macrostates. Binarisation considered 3,228 highly variable genes, identified using a batch-aware mean-dispersion binning strategy [54,98], and ten additional preselected genes of biological interest. Of these, 2,709 received at least one defined assignment. In the resulting abstraction, zero and one represent off- and on-like transcriptomic states, respectively, while ambiguous cases are left unassigned. Expression-density profiles across the annotated populations illustrate the complementary information used by the two methods: single-cell distribution shapes for scBoolSeq, and relative expression shifts between populations for DEA.

To assess whether binarisation preserved macrostate-associated biological programmes, we compared Boolean assignments across process-related gene sets (Fig. 3e; Supplementary Fig. 2). Consistent with expected results, genes involved in cell division, DNA repair, and DNA replication are predominantly assigned to an on-like transcriptomic state in the Rep macrostate, while receiving mostly off-like assignments in granulocyte-like macrostates (Neu, Alt). Neu macrostates show coherent activation of neutrophil-related programmes, with S100a8, S100a9, and Ngp assigned to the on-like state in both conditions and Ly6g receiving an on-like assignment in treated Neu but remaining unassigned in control Neu, whereas these neutrophil-related genes receive a mixture of on- and off-like assignments in the Alt macrostate, indicating that the corresponding programmes are only partially engaged (Supplementary Fig. 1). Importantly, binarisation also captures inter-condition differences: compared with control, treated Neu shows higher proportions of high assignments for both neutrophil-associated and broader inflammatory and immune-related gene sets, suggesting that ATRA further reinforces the neutrophil differentiation programme. This interpretation is consistent with the marked expansion of the Neu population following treatment, from 109 of 5,285 control cells (2.1%) to 806 of 6,855 ATRA-treated cells (11.8%).

#### Model specification

Three major cell fates were identified in the transcriptional landscape: neutrophil-like, alternative differentiated and proliferative states (Neu, Alt and Rep, respectively). Although Cycl displays a distinct cell-cycle programme, it was excluded because the available data did not support an unambiguous dynamical interpretation or placement along the inferred trajectories. All retained terminal configurations were modelled as fixed points. The condition-specific Rep macrostates were initially constrained more generally as trap spaces, but preliminary inference resolved both as fixed points; the final specification therefore imposed fixed-point constraints directly to reduce computational cost. The least differentiated Prom1 macrostates were considered as initial states. Potency scores place Prom2 at an intermediate position between Prom1 and the Neu/Alt populations within the inferred differentiation hierarchy. Accordingly, Prom2 was interpreted as a transitional promyelocytic state preceding terminal differentiation. In contrast, the proliferative Rep state was considered as an alternative fate directly accessible from Prom1, with non-reachability constraints preventing Prom2 from accessing Rep configurations. These interpretations guided the manual specification of intra-condition trajectory constraints. Inter-condition relationships were encoded by preventing condition-specific initial states from reaching differentiated terminal profiles absent from the other condition. In particular, the loss of the alternative differentiation pathway following ATRA treatment motivated a non-reachability constraint preventing the treated Prom1 state from reaching Alt. Finally, universal constraints restricted the fixed points reachable from each initial or intermediate configuration to the corresponding condition-specific terminal profiles (Fig. 3f; Supplementary Table 5).

#### Gene selection

Starting from a regulatory domain comprising 5,198 genes derived from high-confidence mouse DoRothEA interactions (confidence levels A and B), an uncertified soft-stage solution retained 5,016. Solver-certified strong-constant maximisation removed 4,473 components from this propagated domain, leaving 543. Complete-constraint selection subsequently retained 515 genes as a feasible, uncertified incumbent (Supplementary Table 6). This gene set is significantly enriched for genes involved in DNA replication, cellcycle regulation, and transcriptional control (Fig. 4a), consistent with the proliferative regulatory programme identified in the Rep macrostate. Notably, the resulting regulatory domain retains key regulators implicated in PLZF-RAR*α* resistance, including E2F family members (E2f1, E2f2, E2f7 and E2f8) and Ezh2 [77]. The selected domain is also enriched for genes involved in terminal differentiation and regulation of cell population proliferation.

**Fig. 4:**
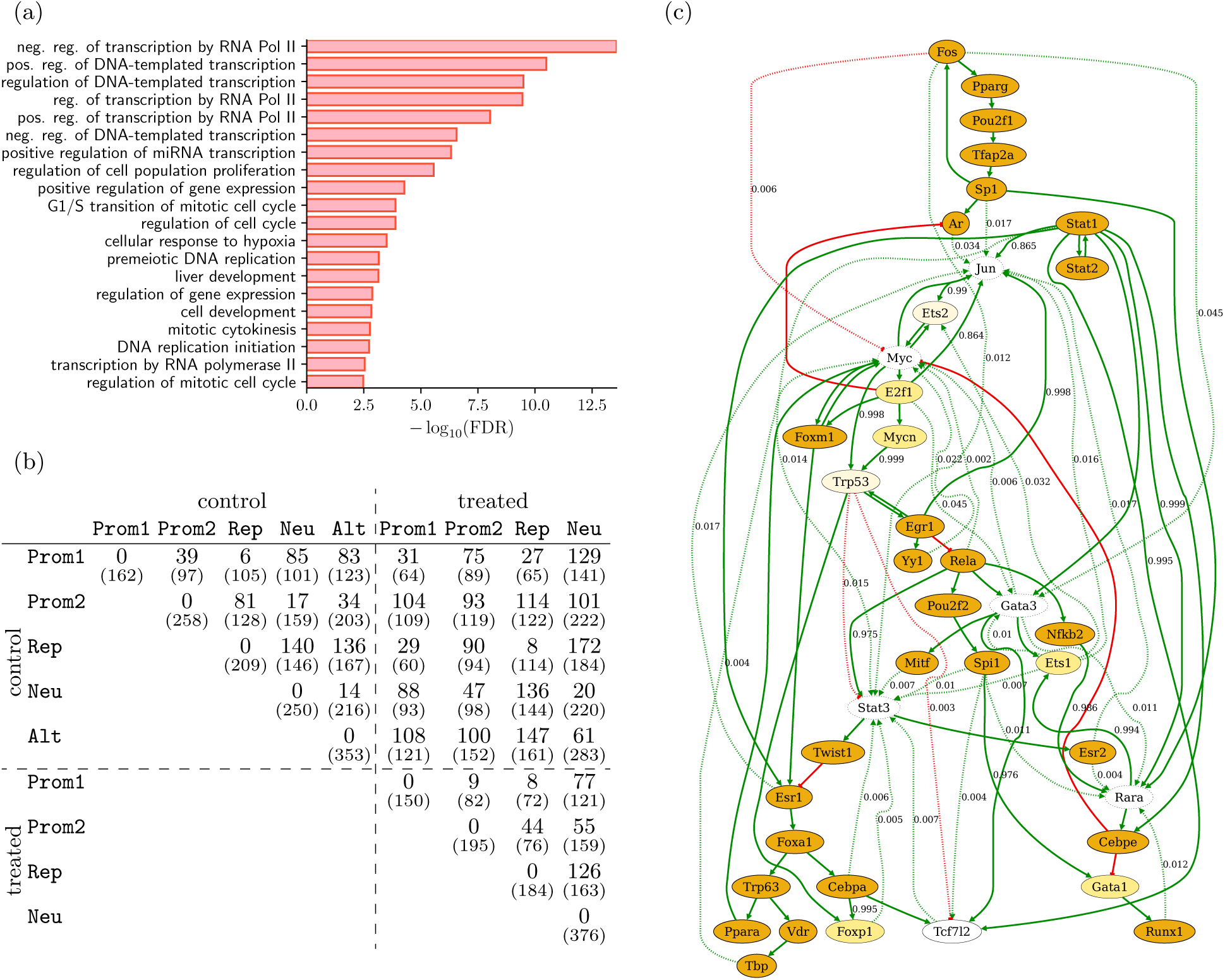
APL logical model analyses. (a) Bar plot of the top 20 over-represented Gene Ontology biological process (BP) terms among the 515 genes identified by BoNesis. The background corresponds to the mouse DoRothEA regulatory domain used for gene selection. Terms are ranked by adjusted p-values. (b) Pairwise distance matrix between macrostates based on the 515 genes selected by BoNesis. Each distance corresponds to the number of genes that are binarised in both macrostates but assigned different values, representing a lower bound on their disagreement. The number in parentheses gives the size of the comparison support, i.e. the number of genes jointly binarised in both macrostates. The corresponding similarity can be obtained by subtracting the distance from this number. (c) Ensemble-aggregated influence graph induced by genes participating in a feedback circuit in at least one of the 1,000 retained BNs. All influences between these genes are retained, not only those directly involved in feedback. Node styles encode the number of distinct inferred logical rules, while edge colours and labels indicate interaction sign and empirical frequency within the retained ensemble, respectively; labels are omitted for interactions present in all 1,000 retained BNs.

To compare the binarised macrostate profiles, we computed pairwise distances over the 515 selected genes (Fig. 4b). Each entry reports a lower bound on Boolean-state differences — the minimum mismatch between two profiles — together with the number of jointly defined genes supporting the comparison. In particular, over their respective comparison supports, the Prom1-Rep and Neu-Alt profiles show strong within-condition similarities, whereas the proliferative Rep profiles remain clearly separated from the differentiated Neu and Alt profiles. Interestingly, although ATRA markedly reshapes the phenotypic composition of the dataset, shared macrostates also exhibit measurable transcriptional differences across conditions, indicating that treatment not only alters cell-state abundances but also remodels their transcriptional profiles.

#### BN inference

An ensemble of 1,000 subset-minimal BNs was inferred from the 515-gene domain under the specified dynamical constraints. As macrostate profiles were partially binarised, inference also completed their undefined values. All inferred BNs contain six minimal trap spaces: five fixed points associated with the specified condition-specific terminal macrostates, and one additional trap space that is unreachable from either inferred Prom1 configuration under the *most permissive* semantics (Supplementary Table 7).

The ensemble-aggregated influence graph induced by feedback-core genes comprises 40 nodes and 91 interactions. Among the retained ensemble, 49 interactions are present in all 1,000 retained BNs, and 27 genes have the same logical rule in all 1,000 retained BNs (Fig. 4c). The complete ensemble graph, with structurally equivalent non-feedback genes collapsed into function families, is provided in Supplementary Fig. 3. The interactions present in all 1,000 retained BNs connect the positive Stat1–Stat2 circuit to a granulocytic-differentiation branch involving Rara and Cebpe [73]. The inhibition of Myc by Cebpe [88] and the positive Myc–E2f1–Foxm1 circuit are likewise present in all 1,000 retained BNs, linking differentiation to regulators of cell-cycle progression and leukemic proliferation [81,66]. These counts describe the retained ensemble and are neither probability estimates nor claims about the complete space of compatible BNs.

#### Comparison with an expert-curated model

We compared the dynamical behaviours of the inferred BNs with those of an expert-curated model of ATRA response in APL [82], hereafter referred to as the comparison model. The comparison focuses on the PLZF-RAR*α* setting, as this model additionally represents wild-type and PML-RAR*α* contexts. In this model, three phenotypes are represented: proliferative, differentiated, and (cell cycle) arrested. Proliferative and arrested phenotypes are reachable in the absence of RA. Upon RA exposure, granulocytic differentiation becomes accessible and the distinct arrested phenotype is lost, while a resistant proliferative fate persists. At the same level of abstraction, a comparable treatment-dependent reorganisation is represented in our inferred BNs. Differentiation and proliferation correspond to the Neu and Rep macrostates, respectively, with both remaining reachable under treatment. The correspondence is less direct for cell-cycle arrest: the Alt macrostate, which exhibits apoptosis-related signatures, likewise becomes unreachable under treatment. Although Alt and cell-cycle arrest represent distinct biological outcomes, both occupy a comparable dynamical role as alternative non-proliferative terminal fates lost upon treatment.

We next examined whether the phenotype-level correspondences identified above are also reflected in the states of shared regulatory components (Fig. 5). As expected, a broad panel of proliferation-associated genes — Ccna2, Ccne2, Cdk4, E2f1, Myc and Skp2 — share the same fate-specific pattern across the comparison model and both scBOLT conditions: they occupy the high state in the comparison model’s proliferative phenotype and both Rep attractors, but the low state in its differentiated and arrested phenotypes and in Neu and Alt. The comparison of non-proliferative fates is more nuanced but equally informative. Alt agrees with the arrested phenotype for 14 of the 18 comparable components, yet is even closer to the differentiated phenotype (15/18). Among jointly defined values, it differs from control Neu only at Cebpe and Runx1. This dual resemblance supports its interpretation as an alternative differentiated state, while explaining why it occupies a dynamical role comparable to that of the arrested phenotype.

**Fig. 5:**
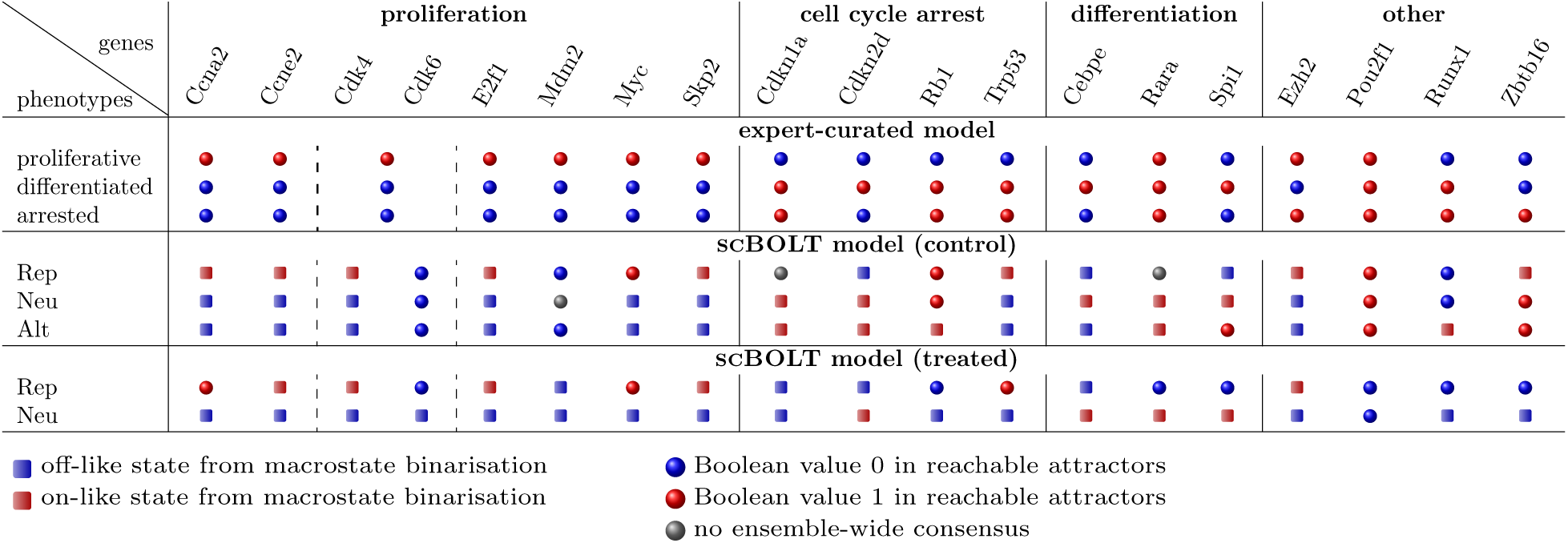
Gene-level assessment of APL attractors. Component values across reachable attractors of the expert-curated model [82] and the 1,000 subset-minimal BNs inferred by scBOLT. Blue and red indicate Boolean values 0 and 1, respectively, while grey indicates no ensemble-wide consensus; squares mark transcriptomic assignments fixed by macrostate binarisation and circles values determined from reachable attractors. For scBOLT, Boolean values 0 and 1 represent off- and on-like transcriptomic states rather than direct protein-activity states. The comparison model’s Cdk4/6 component is shown across the Cdk4 and Cdk6 columns.

Within the differentiation module, Cebpe and Spi1 exactly reproduce the corresponding states of the comparison model in Rep and Neu across both conditions. The remaining differentiation component, Rara, is even more informative, as it reveals a fate-specific distinction absent from the comparison model. In that model, Rara is an exogenous component fixed in the high state, whereas scBOLT models it as an endogenous component governed by an inferred logical rule. Under treatment, Rara retains this state in Neu but adopts the opposite state in the persistent Rep attractor, while no ensemble-wide consensus is recovered for control Rep. Rather than constituting a simple disagreement, this reversal associates the persistent proliferative fate under ATRA with a low Rara state. A second departure concerns Pou2f1: its fixed high state in the comparison model is recovered in every control attractor but reversed in both treated attractors. Together, the conserved modules and these structured departures show that scBOLT recapitulates the principal phenotype-associated patterns of the comparison model while revealing additional fate- and condition-specific distinctions.

#### Phenotype reachability under perturbations

Finally, we evaluated *in silico* all single-gene knock-in and knock-out perturbations of the 40 feedback-core genes to prioritise candidate therapeutic targets and intervention directions according to their effects on phenotype reachability across the BN ensemble (Supplementary Table 10). For each perturbation, a score was computed from lost and preserved attractor reachability. Scores were aggregated over BN–condition pairs; consequently, the same perturbation could produce targeted effects in some pairs and off-target effects in others. The analysis identified ten interventions that render Rep unreachable in all 1,000 retained BNs under both conditions. These include the proliferation-associated regulators Myc, E2f1 and Foxm1, whose roles are already supported by the literature: the PLZF-RAR*α* fusion oncoprotein sustains c-MYC expression in cells surviving RA exposure [81], resistant Rep cells exhibit persistent E2F activity [77], and FOXM1 promotes cell-cycle progression in AML [66]. The Rara→Cebpe┤Myc branch, present in all 1,000 retained BNs, further couples this module to granulocytic differentiation, consistent with C/EBP*ε*-mediated repression of E2F–MYC programmes during cell-cycle exit [88]. Accordingly, fixing Cebpe to the on-like (1) state renders Rep unreachable under both conditions, whereas the corresponding fixation of Rara has this effect only under treatment. Fixing Stat1 or Stat2 to the on-like state instead blocks access to both Rep and Alt in control while preserving Neu. In addition, perturbations affecting only subsets of BNs further expose mechanistic variability and provide model-discriminating experimental predictions.

### 2.3 Case study: hematopoietic differentiation

We applied the scBOLT framework in the context of early blood cell differentiation, using a well-studied dataset built from hematopoietic stem and progenitor cells (HSPCs) in mice [68]. This case study aimed to identify putative regulatory features and logical rules underlying early lymphoid-myeloid lineage commitment. In the original study, individual cells were annotated using FACS index sorting coupled with immunophenotypic classification, spanning major HSPC populations including hematopoietic stem cells (HSCs), multipotent progenitors (MPPs), lymphoid-primed multipotent progenitors (LMPPs), common myeloid progenitors (CMPs), megakaryocyte-erythroid progenitors (MEPs), and granulocyte-monocyte progenitors (GMPs). Additional methodological details are provided in Supplementary Note 2.

Because preliminary differentiation analyses were available from the original STREAM publication [14], scBOLT was applied from the trajectory-inference stage onward, using a preprocessed dataset comprising 1,656 cells and 35,041 unique gene symbols after identifier harmonisation. For latent representation, 2,000 features were selected by LOWESS point-to-curve distance; 100 principal components were computed and the first 40 were retained. Cells were then embedded using spectral embedding [5], a manifold learning approach suited for preserving continuous developmental structures and branching topologies in differentiation landscapes. An elastic principal graph was subsequently inferred from this embedding (Fig. 6a), producing a structure closely matching the original reconstruction and recovering comparable lymphoid and myeloid branching patterns. The reconstructed flat tree was further repurposed for macrostate characterisation by assigning cells to major branching and terminal nodes according to their associated principal graph positions, yielding six macrostates. Macrostate composition analysis (Fig. 6b) shows that S1 is exclusively composed of HSCs, identifying it as the earliest stem-like macrostate.

**Fig. 6:**
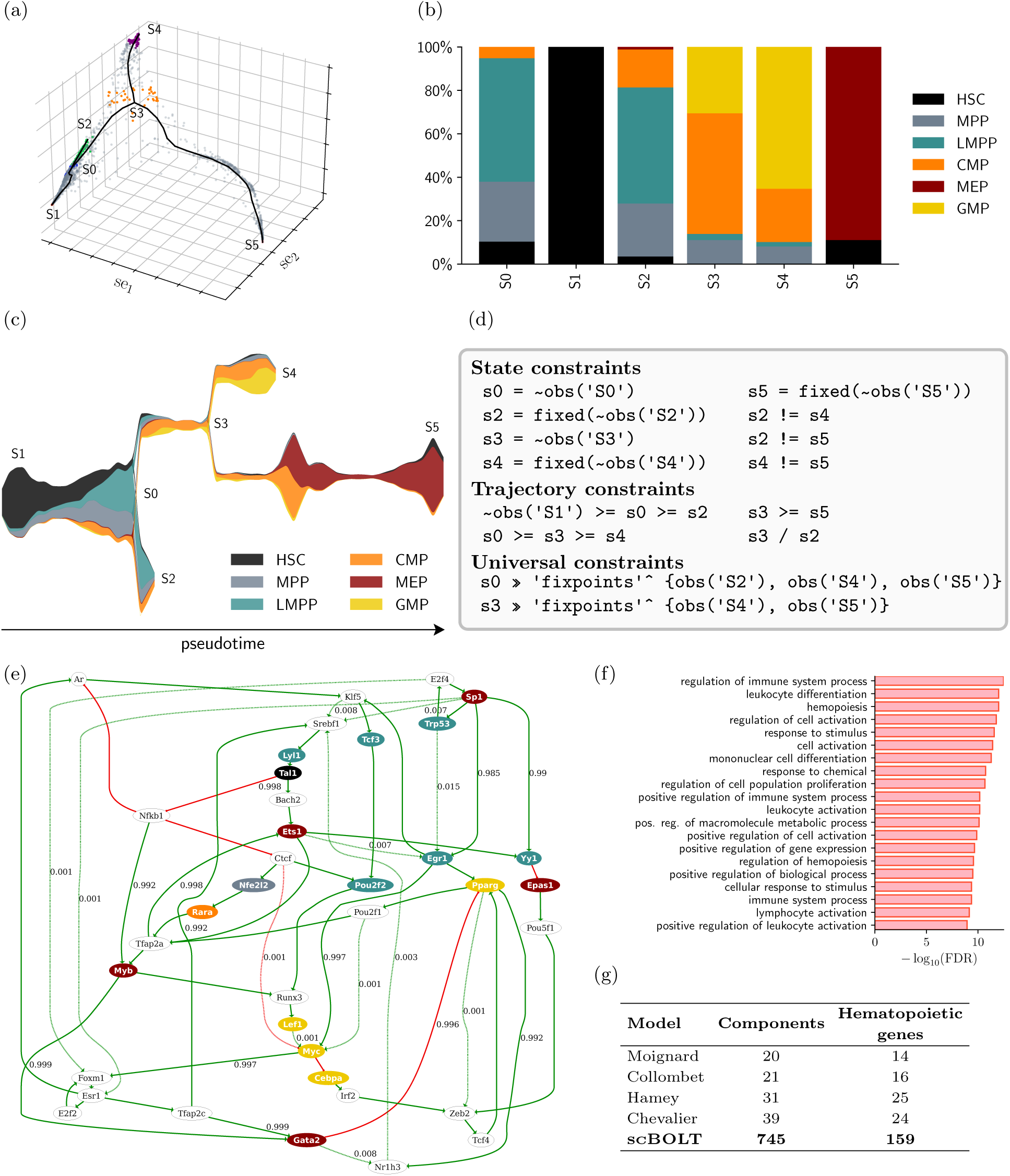
Hematopoiesis analyses. (a) Elastic principal graph defining the major differentiation branches and macrostates. (b) Macrostate composition across hematopoietic populations. (c) Streamplot reconstruction of early differentiation trajectories and lineage bifurcations. (d) BoNesis constraints on states, trajectories and attractors. (e) Ensemble-aggregated feedback core across 1,000 subset-minimal BNs; edge colours encode regulatory signs, labels report empirical selection frequencies within the retained ensemble and are omitted for interactions present in all 1,000 retained BNs, and node colours indicate GO-derived lineage annotations. Lineage colours are defined in panel (b); white denotes unclassified genes. (f) Top 20 GO Biological Process terms enriched among the 745 genes retained by scBOLT relative to the mouse DoRothEA A–C prior, ranked by Benjamini–Hochberg-adjusted *p*-value. (g) Model sizes and numbers of recovered genes from an expanded GO-derived hematopoietic gene set (1,341 genes), combining broad hematopoiesis with stem/progenitor differentiation and major lymphoid and myeloid lineage programmes and their descendant terms (Supplementary Note 2). The 39 components of the published Chevalier model induce a 137-interaction DoRothEA prior [15].

By establishing S1 as the starting point, a streamplot representation was used to reconstruct cell-level early hematopoietic trajectories (Fig. 6c). This dynamic representation reveals an initial transition from the stem-like S1 macrostate toward S0, enriched in multipotent progenitors (MPPs and LMPPs), followed by a bifurcation event leading to either lymphoid or myeloid lineage commitment. The branch from S0 to S2 is associated with progressive lymphoid priming, as S2 remains enriched in MPP and LMPP populations, whereas progression from S0 to S3 marks commitment toward the myeloid compartment, with S3 predominantly composed of CMPs. A second bifurcation subsequently emerges from S3, leading either to S4, predominantly enriched in GMPs, or to S5, which is almost exclusively composed of MEPs, thereby capturing granulocyte-monocyte and erythroid differentiation endpoints, respectively. These results are consistent with the established hematopoietic hierarchy, characterised by progression from HSCs to MPPs, followed by branching toward LMPP and CMP states, the latter further diverging into MEP and GMP trajectories [71,46].

scBOLT subsequently generated Boolean profiles for each macrostate using the consensus procedure, which reconciles scBoolSeq- and DEA-based approaches, the latter corrected using the Benjamini-Hochberg procedure [7]. Among the 4,241 genes considered at this stage, the proportion assigned a Boolean value in each macrostate ranges from 11.5% to 39.3% (Supplementary Table 11), with transitional macrostates S0 and S3 exhibiting comparatively lower binarisation levels, consistent with their intermediate developmental positions. In contrast, terminally differentiated macrostates, particularly S4, display substantially higher binarisation rates, reflecting stronger transcriptional specialisation.

After state abstraction, dynamical constraints were specified by the user in the BoNesis declarative language to restrict model behaviours. Reachability constraints were primarily defined according to the differentiation trajectories reconstructed by the streamplot. Additionally, a non-reachability constraint was imposed between S3 and S2, as no continuous manifold progression directly connects these macrostates without transiting through S0. To restrict long-term dynamical behaviours, universal constraints were additionally imposed on reachable fixed points (Fig. 6d; Supplementary Table 5).

By reconciling dynamical constraints with the mouse DoRothEA prior at confidence levels A–C, scBOLT started from 9,170 candidate genes. The soft stage attained a solver-certified objective vector and retained 9,113 genes, all of which remained compatible after introducing the non-reachability constraint. The strong-constant pass removed 8,367 genes after reaching a solver-certified strong-constant optimum within the propagated domain under the constraints active at that stage, leaving 746 candidates. The complete constraint set retained 745 genes as a feasible, uncertified incumbent (Supplementary Table 6). The final 745-gene domain is significantly enriched for hematopoiesis-associated GO terms relative to the full DoRothEA background, including the GO Biological Process term *hemopoiesis* (GO:0030097; adjusted *p* = 9.89 *×* 10*^−^*^13^; Fig. 6f).

Constrained BN inference subsequently generated 1,000 subset-minimal BNs. Their ensemble-aggregated influence graph contains 697 signed interactions, including 285 present in all 1,000 retained BNs, and its feedback core comprises 37 genes (Fig. 6e). The complete ensemble graph, with structurally equivalent non-feedback genes collapsed into function families, is provided in Supplementary Fig. 4. Each of the 1,000 retained BNs contains three minimal trap spaces, and three attractors are reachable from both S1 and S0 (Supplementary Table 7). The presence of positive circuits within the inferred influence graph is consistent with Thomas’ structural requirement for multistability [91,92,89], providing the regulatory substrate necessary for the imposed double bifurcation, while the dynamical constraints ensure that these circuits are embedded within BN solutions compatible with hematopoietic branching. The feedback core includes factors associated with early hematopoietic commitment and progenitor homeostasis (Tal1 [95], Egr1 [69,60], Trp53 [51]).

Four representative BN frameworks for hematopoietic differentiation have previously been developed: one literature-based model [20], two RT-qPCR-driven models [62,32], and one scRNA-seq-driven model [15]. In contrast to scRNA-seq-based frameworks, RT-qPCR-based approaches require substantial prior biological expertise to predefine regulators of interest during experimental design, inherently restricting inference to targeted low-dimensional regulatory panels. Comparative evaluation across the four frameworks and scBOLT was performed by assessing recovery of an expanded GO-derived gene set covering broad hematopoiesis, hematopoietic stem and progenitor differentiation, and major lymphoid and myeloid lineage programmes (Fig. 6g). Pairwise intersections between their component sets are reported in Supplementary Table 12. While targeted RT-qPCR-based models contain a higher proportion of hematopoiesis-associated genes, scBOLT leverages the transcriptome-wide coverage of scRNA-seq to infer a substantially larger model and recover a greater absolute number of such genes. Furthermore, scBOLT preserves a broader context-aware regulatory landscape: GO terms enriched among the retained genes are dominated by hematopoietic and immune processes, including leukocyte differentiation and cell activation, while also encompassing responses to stimuli and regulation of cell population proliferation. Compared with the previous scRNA-seq-based framework [15], which also relies on the BoNesis engine, scBOLT provides a more automated and modular pipeline. More importantly, the combined use of staged constraint enforcement, clause continuation and domain continuation changed the practical tractability of the full inference problem: when applied to the same 9,170-component input domain, complete dynamical specification and five-clause bound, the historical monolithic gene-selection procedure [15] returned no feasible incumbent within 48 h, whereas scBOLT produced the reported feasible 745-component domain and completed the retained-output workflow in 10 h 10 min (Supplementary Note 3 and Supplementary Tables 8 and 9).

### 2.4 Software implementation

scBOLT is implemented as a lightweight, rule-based software framework for synthesising BNs from multi-condition single-cell transcriptomic data (Fig. 7). It generates intermediate analysis outputs and exports inferred BN ensembles in the standard .bnet format. Host-side execution requires only a Bash interpreter, GNU Make≥ 4.3, and one supported execution backend—Docker, Conda, Mamba, or Micromamba—while module-specific dependencies are provided through controlled environments. The software follows a target-driven dependency paradigm, where each module is defined by its inputs and outputs, and executing a target triggers the recursive construction of dependencies. This design enables the reuse and inspection of intermediate results produced at each module, including data analyses and visualisations, while allowing flexible execution through simple commands (e.g., scbolt <MODULE>). scBOLT further promotes reproducibility, as configuration files explicitly define both software parameters and user-defined modelling choices while sources of non-determinism are controlled by setting random seeds, and flexibility, as users can easily adjust parameters or start the framework from intermediate modules when prior results are available. Exact case-study configuration and notebook paths are listed in Supplementary Table 1, with implementation and performance details in Supplementary Note 3.

**Fig. 7:**
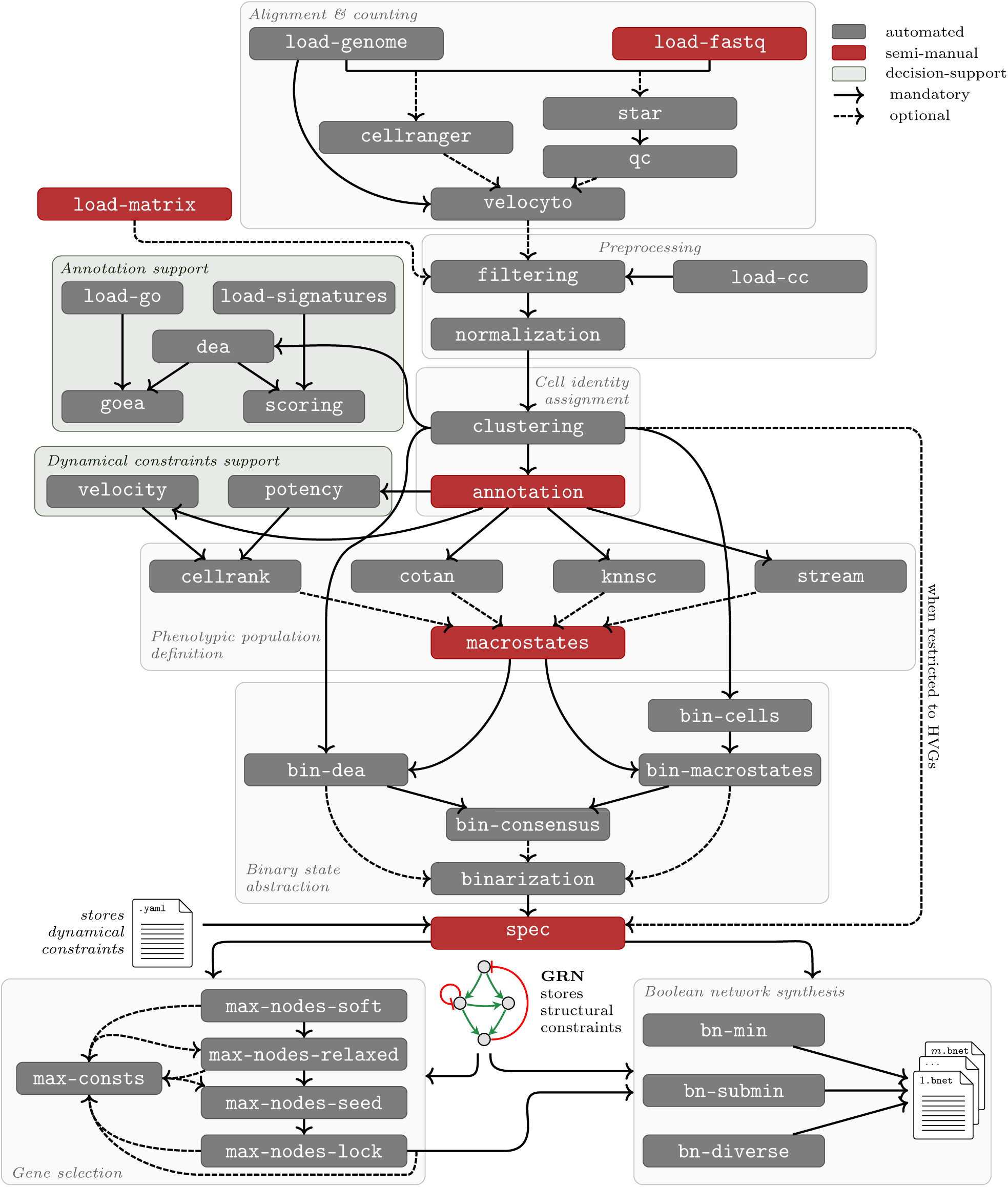
scBOLT implementation. Directed acyclic graph representing the software modules and their dependencies. Boxes denote modules, edges denote their connections, and shaded regions denote higher-level steps. Solid and dashed arrows indicate mandatory and optional paths, respectively. Colours indicate whether modules are automated or semi-manual, while green highlights decision-support steps.

Computational scalability is addressed primarily within the gene-selection stage, which decomposes the full optimisation problem into successive modules with progressively refined regulatory domains and dynamical constraints. An optional strong-constant optimisation pass can be applied after the soft, relaxed or full constraint stage, thereby determining the constraint scope under which strong constants are identified and removed. The multiple optional paths to max-consts in Fig. 7 thus represent alternative scopes for a single pass rather than successive applications. Each selection stage retains a jointly satisfiable component set under its active constraints; omission from such a set does not establish that a component is individually incompatible. Staged constraint enforcement is complemented by clause-bound continuation and domain continuation, which propagate structural witnesses across increasingly complex optimisation problems to guide subsequent searches. Together, these strategies substantially reduce the practical cost of ASP-based solving and make the reported case-study analyses feasible within realistic computational budgets.

scBOLT is designed to be as automated as possible, but some steps remain intentionally semi-manual, allowing users to incorporate expert knowledge and tailor the modelling to the biological processes under investigation and specific biological questions. Semi-automated data retrieval is provided through either load-fastq, which downloads scRNA-seq reads in FASTQ format from SRA/INSDC run accessions, or load-matrix, which downloads precomputed count matrices from GEO Sample accessions (GSM). Other semi-automated modules include annotation for assigning biological identities to cell clusters, macrostates for choosing a method to define phenotypic populations, and spec for specifying dynamical constraints and additional modelling options, including gene prioritisation and the enforced inclusion of specific genes. The software is designed to allow partial execution even when some parameters are undefined, such that failures occur only when reaching targets requiring these parameters, rather than at initialisation. This enables exploration of intermediate steps without enforcing complete configuration upfront. The source code and its documentation are publicly available at https://github.com/bnediction/scbolt.

## 3 Discussion

The construction of mechanistic models has traditionally relied on forward approaches, requiring extensive knowledge of biological macromolecule interactions, as well as iterative refinements to adjust the handmade models. Reverse approaches leveraging omics data offer a promising alternative, enabling modelling directly from observations. However, existing methods remain limited by their difficulty in handling challenging contexts involving unexpected signalling pathways and by their inability to integrate multi-condition datasets. scBOLT addresses these limitations by jointly encoding multi-condition transcriptomic data, user-defined dynamical constraints and prior regulatory knowledge within a unified constraint-based formulation. Its gene-selection strategy, including strong-constant pruning, reduces the search space to improve the tractability of BN synthesis and enable the identification of sparsest BNs consistent with the resulting specification.

Beyond its methodological contributions, scBOLT addresses the need for greater automation by providing modular software that offers a set of strategies for data processing and model construction. Users can therefore select methods suited to their data and biological questions, while interoperable entry points allow analyses to be initiated at different stages and independently generated results to be incorporated. This design preserves user control over biological assumptions and modelling choices, ensuring adaptability across a wide range of biological contexts. Dynamical Boolean constraints remain user-defined; the APL case study provides practical guidance for their formulation in multi-condition datasets, whereas their extension to time-series data remains to be explored.

A central feature of scBOLT is that constraint-driven gene selection becomes an interpretable outcome of BN synthesis. scBOLT first maximises satisfiable nodes, namely components that can be jointly included in a BN satisfying the specification. Nodes not retained may reveal tensions among binarised profiles, dynamical constraints and the prior GRN, thereby guiding targeted revision of these inputs, although their exclusion does not establish that each omitted node is individually incompatible. Among retained nodes, removing strong constants — components admitting a compatible realisation with a constant rule matching all defined configurations — focuses the model on the subset collectively accounting for the state changes represented by the specification, without implying that discarded genes are biologically irrelevant. Edge-minimum and subset-minimal BNs then provide complementary parsimonious explanatory structures whose regulations collectively satisfy the specification. Edge-minimum BNs globally minimise the number of signed interactions. By contrast, subset-minimal BNs need not have minimum cardinality: they require only that no proper subset of their signed interactions supports a compatible realisation. As every compatible BN contains a subset-minimal core, enumerating such cores offers a compact view of irreducible regulatory structures. Diversity-oriented sampling instead favours variation in Boolean clauses and constant assignments among compatible BNs, without guaranteeing either minimum edge cardinality or inclusion minimality.

Nevertheless, computational tractability remains a fundamental challenge across constraint-based BN inference problems, particularly for instances involving large regulatory domains and rich dynamical specifications. Through staged constraint enforcement, clause continuation and adaptive domain continuation, scBOLT extends the feasible scale of constraint-based inference of transcriptome-derived BNs into the several-hundred-component range within practical runtimes. The computationally demanding multi-condition APL case study illustrates this gain: selection of a 515-component domain from 5,198 candidate genes and inference of an ensemble of 1,000 subset-minimal BNs together required just under 24 hours of wall-clock time on a single laptop, comprising 21 hours 21 minutes for gene selection and approximately 2 hours 34 minutes for BN inference. Including upstream analysis, potency scoring and model specification, all logged stages totalled approximately 24 h 8 min (Supplementary Table 8). The hematopoiesis comparison further shows that this combined strategy can make a large, constraint-rich instance accessible within a practical time limit: under the same problem specification, monolithic optimisation returned no feasible solution within 48 h, whereas scBOLT produced the reported 745-component domain in 10 h 10 min. When the strong-constant pass is omitted or applied only after the last active constraint stage, and the complete stage attains the same solver-certified objective vector as the soft stage, that vector is globally lexicographically optimal within the configured input domain and clause bound, matching the guarantee of monolithic BoNesis optimisation [15].

Despite its modularity, flexibility and improved computational tractability, scBOLT remains limited by the scope of its input data. The framework currently focuses on single-cell transcriptomic data and does not support other types of omics data accessible at single-cell resolution such as chromatin accessibility (scATAC-seq) or DNA methylation (snmC-seq). As a result, the reconstructed models may overlook important functional mechanisms acting beyond gene expression. Further improvements could incorporate epigenomic features as additional regulatory layers, which could be facilitated by single-cell sequencing technologies capable of jointly profiling epigenomic and transcriptomic modalities (e.g. 10X Multiome or scNMT-seq [19]), thereby enabling BNs to capture critical inter-omics interactions. In addition, scBOLT does not infer signed dependencies *de novo*, but relies on prior GRNs derived from existing databases, making the quality of inferred models dependent on the completeness of available knowledge.

Future developments of scBOLT could focus on integrating additional regulatory layers to provide a more comprehensive representation of regulatory mechanisms between biological macromolecules. Incorporating GRN inference methods as an alternative to database-derived interactions could help uncover novel molecular relationships. In addition, extending the framework to time-resolved datasets would offer new opportunities to better capture temporal aspects of cellular dynamics. Owing to its modular design, scBOLT can be readily extended by incorporating additional modules within the pipeline, facilitating the integration of alternative strategies for trajectory inference, macrostate characterisation, binarisation, model construction and related steps.

Beyond improving the molecular description of regulatory processes, another promising direction concerns the integration of inferred BNs into higher levels of biological organisation. Current multiscale modelling frameworks assume that intracellular regulatory models are known [61,57,23,49] and simulate how intracellular signalling, cell-cell interactions and microenvironmental processes jointly shape multicellular dynamics. Recent frameworks already support the association of distinct BNs with different cellular populations [48]. An interesting perspective would be to leverage scBOLT as an intracellular BN reconstruction engine for future multiscale modelling frameworks. Combined with spatially resolved datasets, this could support the inference of context-dependent regulatory architectures, opening the way toward multiscale inference frameworks where intracellular BNs are inferred from data rather than assumed a priori.

## 4 Materials and methods

scBOLT provides a semi-automated framework for constructing Boolean network models from multi-condition single-cell transcriptomic data. We first introduce the mathematical formalism underlying Boolean network modelling, followed by a detailed description of the pipeline modules, as illustrated in Figure 7. The implementation relies on custom utilities provided by BoNesisTools, a Python library developed for this work (available at https://github.com/bnediction/bonesistools). scBOLT standardises gene identifiers using organism-specific NCBI gene_info annotations, resolving synonym conflicts by prioritising official nomenclature authority symbols for consistent integration of heterogeneous biological resources.

### 4.1 Boolean Networks

Boolean networks (BNs) are a subclass of executable models [42,90], composed of a finite set of components *i* ε {1*, … , n*}, each of which can be in either Boolean state 0 or 1. For transcriptome-derived scBOLT states, these values denote off- and on-like transcriptomic states, respectively, and not direct protein inactivity or activity. Formally, a BN can be represented as a dynamical Boolean system *f: B^n^→B^n^* where B = {0,1} is the domain, *x* = (*x*_1_*, … , x_n_*) is a configuration of component states, and *f* (*x*) = *f*_1_ (*x*) *, … , f_n_* (*x*) defines the system evolution.

#### State abstraction

A hypercube *h* of dimension *n* is a vector *h* ε {0, 1,*}*^n^*, where each component *i* is either fixed to a Boolean value (*h_i_*ε {0, 1}) or free (*h_i_* = *). A hypercube is an abstraction of a set of configurations: c(*h*) = {*x* ε B*^n^ |V_i_* ε{1*, … , n*}*, h_i_* ε {B→*x_i_* = *h_i_*}. A configuration *x* ε B*^n^* matches a hypercube *h* 0, 1*, ^n^* if the hypercube contains the configuration: *x* ∈ c(*h*). In this work, the binarisation procedure yields partially specified gene activity profiles, which are encoded as hypercubes in {0, 1,*}*^n^*, with fixed components for assigned values and free components for ambiguous cases. These hypercubes represent partially binarised macrostates, and configurations matching them correspond to candidate transcriptomic states.

#### Structural properties

Structural properties of BNs refer to their underlying regulatory structure, which can be partially abstracted through an influence graph (IG) representing regulatory interactions independently of the underlying logical rules. Formally, the IG induced by *f* is a signed directed graph *G*(*f*) = (*V, E*), where *V* = {*v*_1_*, … , v_n_*} is the set of vertices associated with BN components and *E* ⊆ {(*v_i_, v_j_,±*})|*v_i_, v_j_* ε *V,* ± ε{+,–} }is the set of directed edges corresponding to activating and inhibitory interactions. A positive influence from component *i* to component *j* is defined as:

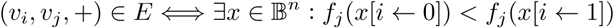

where *x*[*i←b*] denotes the configuration obtained by replacing the *i*-th component of *x* with *b* ε B. Negative influences are defined analogously. In the scBOLT framework, BNs are restricted to locally monotone (unate) Boolean functions, meaning that each regulator acts either as an activator or as an inhibitor, but not both simultaneously:

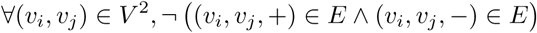

#### Dynamical properties

An updating scheme *µ* of *f* specifies a binary transition relation between configurations →*_µ_* ⊆ B*^n^* ×B*^n^*. Classical update modes include the synchronous semantics, where all components are updated simultaneously, and the fully asynchronous semantics, where only one component is updated at a time. Given an update mode *µ*, a configuration *y* ε B*^n^* is reachable from a configuration *x* ∈ B*^n^*, denoted *x→µ^∗^ y*, if and only if either *x* = *y* or there exists a sequence of transitions *x→_µ_ →_µ_ y*. In this work, dynamical properties are evaluated under the most permissive (MP) semantics [13,74], which is guaranteed to capture all behaviours achievable by any quantitative model consistent with the same logical structure. Intuitively, components may transiently increase or decrease before stabilising, taking an intermediate value (denoted by *) that can be evaluated as active by some components and inactive by others, thereby allowing all compatible interpretations to be explored. Under the MP semantics, any sequence of transitions between two configurations *x, y* ε B*^n^* can be reduced to a direct transition: *x→*_MP_ *→*_MP_ *y* = *x→*_MP_ *y*. The MP semantics facilitates the satisfaction of reachability properties, whereas it makes non-reachability constraints significantly more restrictive.

These notions of reachability naturally lead to the definition of attractors. A non-empty set of configurations *A* ⊆ B*^n^* is an attractor if and only if for each pair of configurations *x, y ε A*, *y* is reachable from *x*, and there is no configuration *z* ε B*^n^*\*A* that is reachable from a configuration in *A*. Within this framework, attractor constraints can be defined by the user, and may represent terminal states, including long-term dynamical behaviours such as cell cycle progression.

Finally, dynamical invariants are defined as sets of configurations that are closed under the system’s evolution. A trap space is a particular class of such invariants, defined as a hypercube *h* such that all associated configurations are mapped to configurations within the same hypercube: *V_x_* ε c(*h*)*, f* (*x*) ε c(*h*). A fixed point is a singleton trap space, i.e. a configuration *x* such that *f* (*x*) = *x*. These notions are used within scBOLT to constrain the model by enforcing trap space or fixed point properties on hypercubes derived from partially binarised macrostates, forming part of the input provided to BoNesis.

### 4.2 Alignment and counting

Raw sequencing data are first processed through standard alignment and counting procedures to obtain gene-level expression matrices using dedicated pipeline functions, including load-fastq, cellranger, and velocyto. scRNA-seq reads are retrieved in FASTQ format from the NCBI Sequence Read Archive (SRA) [47] using parallel-fastq-dump (SRA Toolkit). Reads are then aligned to a reference genome and gene expression is quantified using the cellranger count command (10x Genomics) [98] with default parameters, yielding cell-by-gene count matrices. Spliced and unspliced transcript abundances are estimated from the aligned reads using velocyto run10x [44], providing inputs for downstream analyses of cellular dynamics based on RNA velocity. At this stage, all conditions are processed independently.

### 4.3 Count preprocessing

Count preprocessing steps are performed independently for each condition, using the pipeline functions filtering and normalization, which rely on routines from the BoNesisTools package. Prior to any filtering or normalisation, quality control metrics are computed, including library size, gene detection rate, and the proportion of mitochondrial gene expression, together with cell cycle phase assignment using the supervised learning approach Pairs [83], yielding phase probabilities later used to correct for cell cycle effects. All these measures are computed on raw count matrices to preserve information prior to preprocessing. Genes are then retained based on dropout rate, number of cells in which they are expressed, and total count abundance, while cells are discarded according to gene dropout rate, number of genes detected in each cell, total read count, and proportion of mitochondrial gene expression. Cells with abnormal library sizes are excluded using thresholds derived from the median and a Gaussian-consistent MAD computed on log-transformed total counts. Normalisation is subsequently applied to correct technical variability. Counts are first adjusted by a library size factor to account for differences in sequencing depth, then log-transformed to stabilise variance, and scaled to ensure comparability across genes. Finally, unwanted variation due to cell cycle effects can be mitigated by regressing out log-transformed counts using phase probabilities.

### 4.4 Clustering and data integration

Clustering and data integration are performed to identify shared cell populations across conditions using the pipeline function clustering, with dedicated methods including BBKNN [76], Scanorama [37], or ingest, which operate at different stages of the analysis pipeline to correct for batch effects inherent to sequencing protocols. Principal component analysis is applied to regressed-out counts, optionally restricted to batch-aware highly variable genes (HVGs) identified using either the Cell Ranger mean-binned normalized dispersion strategy [54,98] or the seurat_v3 LOESS-regularized mean-variance strategy [87]. A nearest-neighbour graph is then constructed in the integrated latent space. Cells are subsequently partitioned into clusters using the Leiden algorithm [93]. Transcriptional profiles are finally projected in a low-dimensional embedding, using UMAP [58] or *t*-SNE [53] to approximate the intrinsic manifold structure of the data and support downstream analyses.

### 4.5 Cell annotation

Cells are manually annotated by assigning labels to clusters representing distinct cell populations. To assist the user, DEA is first proposed to identify differentially expressed genes between clusters, by performing Wilcoxon rank-sum tests jointly across conditions on integrated log-normalised counts. A gene is considered overexpressed in a cluster if its log_2_ fold-change is higher than a threshold (default: 0.25) and its corrected *p*-value (default: Bonferroni) does not exceed a predefined threshold (default: 0.05). This cross-condition strategy is justified because the integrated latent space identifies shared phenotypic profiles across conditions. Although strong overrepresentation of one condition within a cluster may confound cluster-level differences with condition-associated sample composition or residual technical effects, this risk is partially mitigated because differential expression compares each cluster against all remaining cells, since the comparison set generally includes other cells from the same condition. Using cluster-level marker genes derived from DEA, GOEA and signature scoring are then proposed using the genes retained by the filtering module as background. The enrichment analysis is performed with GOATOOLS [43], while signature scoring uses hyper-geometric tests to evaluate the overlap between cluster-level marker genes and predefined gene signatures. Signature-associated phenotypes include cell fate [94] and hematopoietic lineage [12], although restricted to mouse datasets.

### 4.6 Trajectory inference

scBOLT provides trajectory inference utilities based on velocity vector estimation and potency scoring. RNA splicing kinetics are inferred using scVelo [8] from spliced and unspliced counts previously estimated with the velocyto module, whereas cell developmental potentials are estimated using CytoTRACE 2 (v1.1.0) [41] from raw gene expression values. To avoid mixing condition-specific transcriptional dynamics, these analyses are performed independently for each experimental group.

### 4.7 Macrostate characterisation

Macrostate characterisation aims to identify subsets of cells representing coherent phenotypic profiles within the transcriptional landscape, each one containing cells originating from a single condition. Several complementary approaches are provided by scBOLT for macrostate inference, including CellRank [45], COTAN [27], STREAM [14], and a novel KNNSC strategy introduced in this work.

First, statistical approaches can be used to infer macrostates. CellRank identifies phenotypic populations using velocity-, potency-, and similarity-based kernels, combined through the generalized Perron cluster cluster analysis (GPCCA) [80,79], whereas COTAN derives phenotypic partitions from pairwise gene co-expression patterns inferred from zero UMI count distributions. In contrast, topology-based approaches characterise macrostates from the geometric structure of low-dimensional embedding spaces. STREAM fits an elastic principal graph to cells embedded in a reduced-dimensional space (default: UMAP) and, although primarily designed for trajectory inference, is repurposed here to derive macrostates by grouping cells mapped to terminal nodes and branching points.

When these approaches do not provide sufficiently discriminative phenotypic populations, scBOLT proposes the *k*-nearest-neighbour-based subclustering (KNNSC) strategy, implemented in the BoNesisTools package, which refines precomputed clusters using local manifold topology. A *k*-nearest-neighbour graph (default *k*: 20) is first estimated with respect to an embedding space (default: UMAP). If multiple connected components are identified in the graph, they are linked through nearest inter-component neighbours in the embedding space to obtain a fully connected graph. Shortest path lengths between cells and clusters’ centroids are then estimated on this graph using Dijkstra’s algorithm to preserve local geometric relationships, and can be interpreted as approximations of geodesic distances on the underlying manifold. Two complementary strategies are proposed for subcluster construction: a centrality-based strategy identifying subsets of cells minimising distances to their own cluster’s barycentre, and a periphery-based strategy identifying subsets of cells maximising distances to other clusters’ barycentres.

### 4.8 Macrostate binarisation

Macrostate binarisation aims to derive coarse-grained transcriptional profiles associated with phenotypic populations. It consists of assigning a partial Boolean vector to each macrostate, where zero and one denote low and high transcriptomic states, respectively, rather than direct protein-activity states. In multicondition settings, binarisation is performed from merged datasets using log-normalised counts, without explicit batch-effect correction. This design preserves transcriptional variability associated with both condition- and macrostate-specific effects while maintaining a unified reference for gene activity states across conditions. When separately profiled biological replicates and replicate-of-origin labels are unavailable, inter-condition differences cannot be reliably disentangled from differences in sample composition or residual technical effects. A globally harmonised binarisation strategy was therefore adopted to ensure a consistent interpretation of Boolean transcriptomic states across all conditions. Three complementary strategies are proposed: scBoolSeq [55], DEA-based binarisation, and consensus-based binarisation.

When using scBoolSeq, the bin-cells module first transforms scRNA-seq pseudocounts into a partial Boolean matrix at the single-cell level, whereas the bin-macrostates module subsequently aggregates these results at the macrostate level through a voting procedure. Optionally, only HVGs can be considered, since genes exhibiting low expression variability across cells (e.g. near-Dirac distributions) provide limited discriminative information for Boolean abstraction. Following the original scBoolSeq methodology, genes are classified into zero-inflated, unimodal or bimodal categories using statistical criteria, leading to distinct binarisation strategies. Genes classified as zero-inflated are binarised using expression thresholding strategies, including non-zero detection or quantile-based thresholding approaches, whereas unimodal genes are binarised using a one-sided statistical test (default *α*: 0.05). For bimodal genes, cells are assigned according to their expression mode.

DEA-based binarisation represents a family of binarisation strategies based on differential expression analyses comparing each macrostate against all other cells from the merged dataset. The bin-dea module supports the Wilcoxon rank-sum test (default), the standard Welch’s *t*-test, and Welch’s *t*-test with overestimated variance to derive partial Boolean abstractions. A gene is considered differentially expressed within a macrostate when its corrected *p*-value (default: Benjamini-Hochberg) is below a predefined threshold (default *α*: 0.05) and its absolute log_2_ fold-change exceeds a threshold (default: 0.5). Genes are subsequently binarised within the corresponding macrostate as up- or down-regulated according to the sign of the associated log_2_ fold-change, whereas non-significant values remain undefined.

Consensus-based binarisation combines scBoolSeq and DEA-based strategies to obtain more stringent binarisation while mitigating unreliable abstraction assignments. A gene within a macrostate receives a low or high transcriptomic assignment when at least one method supports the corresponding state, provided that no contradictory assignment is produced by the other method. This strategy favours conservative assignments while preserving complementary information captured by each method. Note that the scBoolSeq methodology may lead to misleading binarisation results under high dropout rates, potentially classifying genes as zero-inflated due to technical rather than biological dropouts. Consequently, low assignments derived from zero-inflated profiles do not override contradictory high assignments inferred by DEA-based binarisation.

### 4.9 Model specification

Following state abstraction, partially binarised macrostates are used to specify dynamical constraints through the declarative BoNesis language [16], describing expected cellular behaviours. This step remains largely manual, allowing users to tailor biological assumptions and modelling objectives to their specific experimental context, whereas the spec module verifies the syntactic correctness of the resulting BoNesis specifications. Users first define observations (encoded as hypercubes) and configurations (encoded as Boolean vectors) matching macrostates. A broad range of dynamical properties can then be specified, including (i) configuration restrictions such as equality or inequality, (ii) attractor-related properties, such as forcing configurations to be fixed points or enforcing the existence of trap spaces in which observations are fixed, (iii) reachability and non-reachability relationships under Most Permissive semantics, and (iv) universal properties such as the reachability of all fixed points from a given configuration.

In cross-sectional multi-condition settings, scBOLT recommends specifying dynamical constraints independently for each condition, guided by condition-specific trajectory inference analyses. Additional constraints should be introduced to enforce inter-condition inequalities between configurations and inter-condition non-reachability relationships between configurations, guided by the non-juxtaposition of condition-specific regions within integrated embedding spaces. This guideline is motivated by the absence of separately profiled biological replicates and replicate-of-origin labels, which prevents reliable disentanglement of inter-condition biological variability from inter-individual or residual technical effects.

This recommendation assumes that each condition captures the complete biological process under study. When conditions instead correspond to complementary snapshots or successive stages of a shared biological trajectory (e.g. distinct differentiation stages sampled independently), inter-condition reachability constraints may remain biologically meaningful. However, such settings require particular caution, as residual batch or inter-individual effects may still induce spurious dynamical relationships.

### 4.10 Gene selection

Prior to BN inference, scBOLT reduces the prior regulatory domain through a sequence of optimisation problems implemented with BoNesis [17,18,16]. For a given domain, gene selection maximises the number of satisfiable nodes, defined as components that can be jointly included in at least one BN satisfying the active dynamical constraints for some completion of the partially defined configurations. When specified, user-designated important nodes define the higher-priority component of a lexicographic objective: the solver first maximises the number of retained important nodes and then the total number of retained nodes. Certification therefore applies to this objective vector and does not, by itself, establish an unconstrained maximum of its lower-priority second component. Mandatory nodes are required in every solution.

Satisfiable-node maximisation is performed through staged constraint enforcement. Dynamical constraints are divided into three classes according to their computational complexity: (i) soft constraints, comprising configuration equality and inequality, reachability and attractor-related properties; (ii) intermediate constraints, comprising non-reachability and final non-reachability properties; and (iii) hard constraints, comprising universal properties over fixed points or reachable attractors. Four node-selection modules, each operating under an independently configurable solving-time budget, successively optimise the satisfiable-node objective: (i) max-nodes-soft under soft constraints; (ii) max-nodes-relaxed after introducing intermediate constraints; (iii) max-nodes-seed, performing an initial bounded-time optimisation under the complete constraint set; and (iv) max-nodes-lock, which resumes optimisation from an uncertified solution previously obtained under the complete constraint set, making its retained nodes mandatory and searching for additional satisfiable nodes within the domain retained before hard constraints were introduced. When a module introduces no additional constraint class, it forwards the preceding domain and structural witness without launching a new optimisation. In both case studies, the lock stage was invoked with a zero optimisation budget; it performed no additional search and propagated the seed-stage incumbent (Supplementary Note 3 and Supplementary Table 6).

A single optional max-consts pass identifies and subsequently removes a maximum-cardinality set of components that are jointly strong constants within the same compatible joint realisation of a BN and its configuration completions. A component *g* is a strong constant with value *v* ε {0, 1} in such a realisation when *f_g_*≡ *v* and every completed configuration assigns *fg* = *v*. Strong constancy is therefore evaluated within a compatible realisation rather than universally across all compatible BNs. The pass may use the soft, relaxed, or full constraint scope, or be disabled altogether. Strong constants are evaluated under the constraints active at the selected scope, and the reduced domain is passed to the following module or to BN inference. When several optimal realisations exist, only one maximum-cardinality set is removed; alternative optima may identify different sets. Consequently, components retained in the reduced domain may still be strong constants in subsequently inferred BNs. BNs inferred over this reduced domain are termed *sparsest BNs* with respect to the strong-constant criterion. This component-level sparsity is distinct from edge sparsity.

To improve tractability, scBOLT provides two continuation strategies: clause continuation solves a sequence of problems allowing *q* = 1*, … , q*_max_ clauses per Boolean function, passing the retained structural witness from one bound to the next as a Clingo heuristic, whereas domain continuation searches adaptive candidate subdomains in parallel to acquire a witness and uses it to guide expansion towards the complete input domain, where a final solver portfolio attempts optimality certification. When both strategies are enabled, domain continuation operates within each successive clause-bound problem. Intermediate clause bounds and domain waves are controlled by separate patience limits within the stage-specific solving-time budget. Both strategies can be enabled independently for each of the four node-selection modules, but do not apply to max-consts.

Optimality certification applies to the exact input domain, constraint set, clause bound and configured objective vector considered by each module. When optimisation is interrupted, the best retained domain and witness are recorded as an uncertified incumbent. Because only one domain is propagated when several optima are tied, downstream optimisations remain conditional on the domain selected upstream. When the strong-constant pass is omitted or applied only after the last active constraint stage, and the complete stage attains the same solver-certified objective vector as the soft stage, that vector is globally lexicographically optimal within the configured input domain and clause bound. When enabled, strong-constant filtering remains optimised within the particular propagated domain and does not establish a joint optimum across tied upstream domains.

### 4.11 BN inference

Using the BoNesis framework, BN inference jointly synthesises BNs and Boolean configurations matching the macrostate-associated hypercubes while satisfying the user-defined dynamical constraints *C* within the regulatory subgraph *G* retained after gene selection:

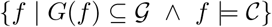

Each inferred BN is returned with a compatible completion of the macrostate-associated hypercubes. scBOLT provides three complementary inference modules for exploring the solution space: (i) bn-min, which enumerates edge-minimum BNs, i.e. compatible BNs with the minimum possible number of regulatory interactions; (ii) bn-submin, which enumerates BNs associated with subset-minimal signed influence graphs, such that no proper subset of their signed interactions supports a compatible realisation; and (iii) bn-diverse, which samples distinct compatible BNs using heuristics that favour diversity in Boolean clauses and constant assignments across retained solutions, without guaranteeing either minimum edge cardinality or inclusion minimality. bn-submin thus explores irreducible regulatory structures, whereas bn-diverse explores logical variability among compatible BNs.

## Code and data availability

The scBOLT software is available at https://github.com/bnediction/scbolt (archived at Software Heritage swh:1:dir:a3a02dc30fe79d996e238c35b2ece3b2a7515a38). Data, scripts, and notebooks that produced the results of the case studies are available at https://github.com/bnediction/scbolt-case-studies (archived at https://doi.org/10.5281/zenodo.22959149). They employed scBOLT version 1.0.0 (swh:1:rev:b470a8c0201d02d17e5024cd833e7b6e4acf957b).

## Acknowledgements

We are grateful to Estelle Duprez and her team for insightful discussions and for their help in interpreting the results of our analyses performed on their single-cell APL dataset.

This research was primarily supported by France 2030 project “AI4scMED” operated by ANR (grant number ANR-22-PESN-0002). LP also acknowledges support from ANR in the scope of the project BNeDiction (grant number: ANR-20-CE45-0001) and RD2BOOL (grant number: ANR-23-CE45-0014).

## Notes

https://github.com/bnediction/scbolt

https://github.com/bnediction/scbolt-case-studies

### Competing Interest Statement

The authors have declared no competing interest.

