## Supplementary material for "Scalable and semi-automated Boolean network inference from multi-condition single-cell transcriptomes with scBOLT"

#### Supplementary Notes

Throughout this document, Boolean values 0 and 1 in transcriptome-derived abstractions denote off- and on-like transcriptomic states, respectively, rather than direct protein inactivity or activity.

##### Supplementary Note 1: Additional methodological details for the APL case study

**Dataset and experimental design.** The APL analysis used the single-cell RNA-sequencing dataset GSE181190 from Poplineau *et al.* [1], comprising an untreated control sample (GSM5492245) and an ATRA-treated sample (GSM5492246) of leukemic cells recovered from recipient mice transplanted with PLZF–RAR$\alpha$ cells; ATRA was administered in vivo for seven days in the treated arm. Each condition was represented by a single 10x library. GEO metadata report cohorts of three to five recipient mice for both experimental arms, but individual mouse identities were not available in the deposited data. The GEO matrices initially contain 5,786 control and 7,416 treated cells. Filtering used MAD-deviation thresholds on log-transformed library sizes: cells falling more than three Gaussian-consistent median absolute deviations (MADs) below the median or more than two MADs above it were excluded. Cells for which mitochondrial transcripts accounted for more than 30% of total counts were also excluded. This retained 5,285 control and 6,855 treated cells. For the latent representation, 2,000 highly variable genes were selected by LOESS before computing 15 principal components. The analysis used a 14-neighbour integrated graph, a Leiden resolution of 0.4, and a UMAP embedding with minimum distance 0.5 and spread 2.0.

**Cell annotation.** Manual annotation was supported by cluster-level differential-expression, GO Biological Process enrichment and signature-scoring analyses of the integrated dataset. The corresponding result tables are provided under `apl/results/annotation/`: `dea/markers.csv`, `goea/basic.xlsx` and `scoring/signature_pvals_adj.csv`.

**Macrostate construction and binarisation.** Six cell populations were annotated as `Prom1`, `Prom2`, `Rep`, `Cycl`, `Neu` and `Alt`. The $k$-nearest-neighbour-based subclustering (KNNSC) method then defined nine condition-specific macrostates of 100 cells each. Central subsets of `Prom1` and `Prom2` and peripheral subsets of `Rep` and `Neu` were retained in both conditions; a peripheral `Alt` macrostate was additionally retained in control. `Cycl` cells were therefore represented in the integrated annotation but did not define a macrostate.

For binarisation, an independent binning-based feature-selection step used 20 expression bins and no fixed top-gene cutoff, identifying 3,228 highly variable genes. The configuration additionally requested 37 genes of biological interest; 27 were already among the selected genes and the remaining 10 were added explicitly, yielding 3,238 genes for binarisation. Consensus binarisation then combined scBoolSeq [2] and differential-expression assignments. Differential expression used a Benjamini–Hochberg-adjusted significance threshold of 0.05 and an absolute $\log_2$ fold-change threshold of 0.5. For genes classified as zero-inflated by scBoolSeq, zero expression values were left undefined rather than automatically assigned to the low (0) transcriptomic state. Of the 13,239 genes shared after preprocessing, 2,709 received at least one defined consensus assignment across the nine macrostates and entered model specification.

**Dynamical specification and model construction.** The complete human-readable dynamical specification is summarised in Supplementary Table 5, while its executable version is provided in the companion repository. The prior regulatory domain was the mouse DoRothEA network restricted to confidence levels A and B. The soft-constraint stage retained 5,016 of 5,198 candidates as an uncertified incumbent. The strong-constant pass reached a solver-certified strong-constant optimum within the propagated 5,016-gene domain under the constraints active at that stage and removed 4,473 components, leaving 543. All 543 remained compatible after introducing non-reachability constraints, whereas the complete constraint set retained a feasible 515-gene incumbent whose final cardinality was not solver-certified. Boolean functions were limited to four clauses, and the random seed was 10. Clause and domain continuation were used during node selection. The lock stage was invoked with a zero optimisation budget; it performed no additional search and propagated the seed-stage incumbent. Finally, 1,000 subset-minimal BNs were enumerated.

For ensemble visualisation, the signed influence graphs of these 1,000 BNs were merged, and each empirical edge frequency was defined as the fraction of retained BNs containing that signed influence. In the full graph, non-feedback genes with the same signed predecessor and successor patterns in the same edge-frequency bins were collapsed into composite function-family nodes, while feedback genes were kept separate (Supplementary Fig. 3). Node shading represents the empirical frequency of the most common DNF implicant structure for each gene, and edge labels and line styles represent empirical interaction frequency within the retained ensemble. A frequency of one means that the feature is present in all 1,000 retained BNs; these frequencies are neither probabilities nor claims of universality over the complete space of compatible BNs.

#### Supplementary Note 2: Additional methodological details for the hematopoiesis case study

**Dataset and experimental design.** The analysis used the mouse HSPC dataset GSE81682 [3]. It comprises 1,656 cells spanning HSC ($n = 375$), MPP ($n = 234$), LMPP ($n = 246$), CMP ($n = 325$), MEP ($n = 356$) and GMP ($n = 120$) annotations. The published normalised expression matrix was stored as the primary matrix, while raw HTSeq counts matched to the same cells and genes were retained in a separate count layer. Broad cell-type labels were reconstructed from the published immunophenotypic annotations and cluster assignments. Cells expressing fewer than 100 features and genes detected in fewer than five cells above the published expression cutoff of 1 were excluded. Identifier harmonisation reduced 35,077 input features to 35,041 unique gene symbols. For the latent representation, 2,000 features were selected using LOWESS point-to-curve distance (span 0.01); 100 principal components were computed and the first 40 were retained. A four-dimensional spectral embedding was constructed from a 15-neighbour fuzzy graph.

**Macrostate construction and binarisation.** An elastic principal graph was fitted to the spectral representation using STREAM [4], with 20 $k$-means centroids used only to initialise the graph, $\alpha = 0.01$, $\mu = 0.02$, $\lambda = 0.01$, branch extension parameter 0.8 and no pruning. Six macrostates were obtained: S0 ($n = 56$), S1 ($n = 26$), S2 ($n = 96$), S3 ($n = 46$), S4 ($n = 31$) and S5 ($n = 47$). S1 represents the HSC root; S0 and S2 are enriched in multipotent and lymphoid-primed progenitors; S3 represents the CMP branch; and S4 and S5 represent GMP and MEP endpoints, respectively.

For binarisation, a separate feature-selection step used the same binning method with 20 expression bins and no fixed top-gene cutoff, selecting 4,241 genes. Consensus binarisation then used the same scBoolSeq and differential-expression criteria as in the APL analysis, without an additional forced-gene list. In this analysis, zero expression values for genes classified as zero-inflated by scBoolSeq were directly assigned to the low (0) transcriptomic state. After removal of genes undefined in all six macrostates, 2,396 genes entered model specification.

**Dynamical specification and model construction.** The reconstructed differentiation tree imposed the reachability constraints S1 $\rightarrow$ S0 $\rightarrow$ S2, S0 $\rightarrow$ S3 $\rightarrow$ S4, and S3 $\rightarrow$ S5. S2, S4 and S5 were fixed points, and direct reachability from S3 to S2 was forbidden. The mouse DoRothEA A–C prior contained 9,170 candidate genes. The soft stage attained a solver-certified objective vector and retained 9,113 genes, unchanged after adding non-reachability constraints. The strong-constant pass reached a solver-certified strong-constant optimum within the propagated 9,113-gene domain under the constraints active at that stage and reduced the domain to 746 genes. The complete constraint set retained a feasible, uncertified 745-gene incumbent. The lock stage was invoked with a zero optimisation budget; it performed no additional search and propagated the seed-stage incumbent. Selection used at most five clauses per Boolean function, a random seed of 10, and clause and domain continuation from the relaxed stage. It yielded 1,000 subset-minimal BNs.

**GO-derived hematopoietic reference set.** For cross-model comparison, an expanded reference set was constructed from 30 GO Biological Process roots grouped as follows: general hematopoiesis (`GO:1903706`, `GO:0030097`); HSC-associated processes (`GO:0060218`, `GO:1902036`, `GO:1902033`, `GO:0035162`); MPP-associated processes (`GO:0002244`, `GO:1901532`); LMPP-associated processes (`GO:0002320`, `GO:0030098`, `GO:0046650`, `GO:0046649`, `GO:0002521`); CMP-associated processes (`GO:0002318`, `GO:0030099`, `GO:0061515`, `GO:0045637`); MEP-associated processes (`GO:0030219`, `GO:0035855`, `GO:0045652`, `GO:0030218`, `GO:0048821`, `GO:0045646`); and GMP-associated processes (`GO:0036230`, `GO:0071621`, `GO:0030851`, `GO:0030224`, `GO:0042117`, `GO:0002548`, `GO:0045655`). Each root was expanded to all descendant terms in the GO directed acyclic graph, and mouse genes annotated to any resulting term in the archived NCBI `gene2go` resource were pooled after conversion to gene symbols. The resulting union comprises 1,341 unique genes.

The executable construction is provided in `hematopoiesis/notebooks/bn.ipynb`.

#### Supplementary Note 3: Computational implementation, reproducibility and performance

Exact configuration files, command lines, intermediate outputs and figure-generation notebooks are provided in the companion repository https://github.com/bnediction/scbolt-case-studies. The results reported in the manuscript are archived at https://doi.org/10.5281/zenodo.22959149 and were generated with scBOLT 1.0.0, BoNesisTools 1.7.0, BoNesis 0.7.0, Clingo 5.8.0.

**Numerical reproducibility qualification.** Within the reference Intel Meteor Lake environment, repeated runs produced strictly identical outputs for all tested single-cell analysis stages under the same pinned software environment and controlled numerical settings. Cross-architecture identity was evaluated only for the numerical outputs covered by the BoNesisTools golden suite and for DEA-based binarisation. Matching SHA-256 digests were obtained on qualified Linux x86-64 environments spanning Intel Meteor Lake (`meteorlake`) and AMD Zen 3 (`znver3`). For the golden suite, the combined `GOLDEN_NUMERICAL_FAMILY_SHA256` digest incorporates the logical names, shapes, data types and exact bytes of the numerical arrays recorded for every covered stage and diagnostic checkpoint. Outputs produced by `pypairs`, scVelo, CellRank, STREAM and scBoolSeq were not included in this cross-architecture qualification. Consensus profiles incorporating scBoolSeq assignments were therefore also outside its scope. Their strict numerical identity across microarchitectures remains undetermined, and no strict output-identity claim is made for unlisted microarchitectures.

**Scope of BN solver reproducibility.** BN optimisation provides a distinct guarantee. When the complete configured input domain and final clause bound are optimised directly, without domain or clause continuation, completed Clingo optimisation certifies the globally optimal lexicographic value of the configured objective vector. It does not establish uniqueness of the retained component set, because several sets may be tied at that optimum. With fixed software, inputs, seed, solver options and single-threaded execution, repeated runs on one machine can be operationally repeatable, but certification alone does not guarantee identical selected components. Time-limited adaptive or continuation-based optimisation may instead return hardware- and schedule-dependent incumbents.

The staged APL and hematopoiesis analyses were run under Ubuntu 22.04.5 LTS on a laptop equipped with an Intel Core Ultra 7 155H processor and 64 GB of RAM. For both analyses, the omics workflow used 16 parallel jobs, and a 50 GB memory limit was enforced during gene selection and BN inference. Reported runtimes for these staged runs pertain to this host; time-limited incumbents may vary with hardware and solver scheduling.

All satisfiable-node optimisation problems were solved using a single Clingo thread per solve. This was the configuration historically selected and used by the authors for satisfiable-node maximisation. The choice was based on empirical benchmarks in which single-threaded Clingo search outperformed multi-threaded search in the large majority of tested instances for this objective. During domain continuation, the `jobs` parameter instead controls the parallel execution of up to 16 independent single-threaded subproblems and is not the number of threads assigned to one solver instance. Multi-threaded solving was used only for BN enumeration, for which it improved solution throughput (Supplementary Table 9).

For APL, the logged upstream workflow from matrix loading through consensus macrostate binarisation required 10 min 55 s. Potency-score computation for both conditions required 2 min 4 s. Gene selection comprised a 15 h soft-stage timeout, 52 s for strong-constant maximisation, 1 h 20 min 43 s for relaxed node maximisation and a 5 h seed-stage timeout. Enumeration of 1,000 subset-minimal BNs required approximately 2 h 34 min, corresponding to approximately 24 h 8 min of cumulative computation across these stages.

For hematopoiesis, the retained logs record 42 s for STREAM macrostate construction, 38 s for binarisation and 16 s for model specification. Gene selection was resumed across multiple runs: soft and relaxed node maximisation required 5 min 48 s and 1 h 13 min 42 s, respectively; strong-constant maximisation required 1 h 13 min 5 s; and seed refinement ran for 7 h 29 min 14 s before being killed by the operating system after reaching the memory limit. In both case studies, the lock stage was invoked with a zero optimisation budget; it performed no additional search and propagated the seed-stage incumbent. The retained-output selection stages therefore totalled approximately 10 h 2 min. Enumerating 1,000 subset-minimal BNs required 6 min 9 s. Pre-STREAM preprocessing was not logged, and the stages ran separately, so no comparable end-to-end hematopoiesis runtime is reported.

**Historical monolithic baseline.** To evaluate the combined practical gain from staged constraint enforcement, clause continuation and domain continuation, the historical joint max-nodes/max-strong-constants procedure was evaluated with a 48 h time limit. The run used the same complete dynamical specification, 2,396 data nodes and full modern mouse DoRothEA A–C domain comprising 9,170 nodes and 32,473 edges. Boolean functions were canonical and limited to five clauses. Clingo used optimisation mode with the `usc` strategy and the lexicographic objective of first maximising satisfiable nodes and then maximising strong constants. The run was launched with `JOBS=1 ./scripts/run_historical_inference_algorithm.sh`. The explicit `JOBS=1` environment override took precedence over the `jobs: 16` value in `config/params-abc.yml`; the wrapper therefore passed `-jobs 1`, and Clingo used one internal thread. Unlike the scBOLT run, the baseline addressed the complete domain and final clause bound directly, without staged constraint enforcement, clause continuation or domain continuation. The 48 h time limit was reached with no incumbent found. Thus, the baseline used the authors' historically preferred configuration for this task, rather than an intentionally restrictive setting.

### Supplementary Tables

**Supplementary Table 1:** Configuration provenance for the two case studies. Paths are relative to the root of the companion repository.

| Case study | Workflow configuration | Dynamical specification | Analysis notebooks |
|---|---|---|---|
| APL | `apl/params-gsm.yml` | `apl/spec.yml` | `apl/notebooks/omics.ipynb`; `apl/notebooks/bn.ipynb` |
| Hematopoiesis | `hematopoiesis/config/params-abc.yml` | `hematopoiesis/config/spec.yml` | `hematopoiesis/notebooks/omics.ipynb`; `hematopoiesis/notebooks/bn.ipynb` |

**Supplementary Table 2:** Datasets and experimental designs used in the two case studies.

| Case study | Accession | Biological material | Conditions | Retained cells |
|---|---|---|---|---|
| APL | GSE181190 | Leukemic cells from PLZF–RAR$\alpha$ transplant recipients | Control; 7-day in vivo ATRA treatment | 5,285 control; 6,855 treated |
| Hematopoiesis | GSE81682 | Mouse hematopoietic stem and progenitor cells | Single condition | 1,656 cells |

**Supplementary Table 3:** Principal preprocessing, macrostate and binarisation settings. Values not listed here retain their scBOLT defaults and are recorded in the complete configuration files identified in Supplementary Table 1.

| Item | APL | Hematopoiesis |
|---|---|---|
| Input after filtering | 5,285 control and 6,855 treated cells; 13,239 common genes | 1,656 cells; 35,041 genes after symbol harmonisation |
| Quality-control thresholds | Log-library size between median $-3$ MADs and median $+2$ MADs; mitochondrial counts at most 30% of total counts | Published expression cutoff 1; at least 100 features per cell and five cells per gene |
| Omics HVG selection | LOESS; top 2,000 HVGs | LOWESS point-to-curve distance; top 2,000 features; span 0.01 |
| Latent representation | 15 PCs; 14-neighbour graph; UMAP minimum distance 0.5 and spread 2.0 | 40 of 100 PCs; four-dimensional spectral embedding from a 15-neighbour graph |
| Macrostate construction | KNNSC; 100 cells per macrostate; nine condition-specific macrostates | STREAM; graph initialisation from 20 $k$-means centroids; $\alpha = 0.01$, $\mu = 0.02$, $\lambda = 0.01$; extension 0.8; no pruning |
| Binarisation HVG selection | Binning; 20 expression bins; no fixed top-HVG cutoff; 3,228 selected genes plus 10 additions from a 37-gene list | Binning; 20 expression bins; no fixed top-HVG cutoff; 4,241 selected genes; no requested-gene list |
| Consensus binarisation | scBoolSeq and DEA consensus; zero values left undefined for zero-inflated genes | scBoolSeq and DEA consensus; zero values assigned to the low (0) state for zero-inflated genes |
| DEA thresholds | Benjamini–Hochberg $\alpha = 0.05$; $\lvert \log_2 \mathrm{FC} \rvert \geq 0.5$ | Benjamini–Hochberg $\alpha = 0.05$; $\lvert \log_2 \mathrm{FC} \rvert \geq 0.5$ |
| Genes defined in at least one macrostate | 2,709 | 2,396 |

**Supplementary Table 4:** Cell-population composition after quality-control filtering. Counts are followed by percentages calculated within each APL experimental condition and across the full hematopoiesis dataset. Percentages may not sum to exactly 100% because of rounding.

| APL | | | Hematopoiesis | |
|---|---|---|---|---|
| Annotation | Control, $n$ (%) | ATRA-treated, $n$ (%) | Annotation | $n$ (%) |
| `Prom1` | 1,534 (29.0) | 2,281 (33.3) | HSC | 375 (22.6) |
| `Prom2` | 2,061 (39.0) | 1,221 (17.8) | MPP | 234 (14.1) |
| `Rep` | 923 (17.5) | 1,261 (18.4) | LMPP | 246 (14.9) |
| `Cycl` | 408 (7.7) | 1,234 (18.0) | CMP | 325 (19.6) |
| `Neu` | 109 (2.1) | 806 (11.8) | MEP | 356 (21.5) |
| `Alt` | 250 (4.7) | 52 (0.8) | GMP | 120 (7.2) |
| Total | 5,285 (100.0) | 6,855 (100.0) | Total | 1,656 (100.0) |

**Supplementary Table 5:** Human-readable summary of the dynamical specifications. Complete machine-readable specifications are provided in the companion repository.

| Constraint class | Configurations involved | Biological interpretation |
|---|---|---|
| | **APL** | |
| Fixed points | Control: `Rep`, `Neu`, `Alt`<br>Treated: `Rep`, `Neu` | Terminal macrostates represented as stable phenotypes. |
| Reachability | Control:<br>– `Prom1` $\rightarrow$ `Rep`<br>– `Prom1` $\rightarrow$ `Prom2` $\rightarrow$ `Neu`<br>– `Prom2` $\rightarrow$ `Alt`<br>Treated:<br>– `Prom1` $\rightarrow$ `Rep`<br>– `Prom1` $\rightarrow$ `Prom2` $\rightarrow$ `Neu` | Proliferative and differentiation trajectories supported by the data. |
| Non-reachability | Control `Prom2` $\not\rightarrow$ control `Rep`<br>Control `Prom2` $\not\rightarrow$ treated `Rep`<br>Control `Prom1` $\not\rightarrow$ treated `Neu`<br>Treated `Prom2` $\not\rightarrow$ control `Rep`<br>Treated `Prom2` $\not\rightarrow$ treated `Rep`<br>Treated `Prom1` $\not\rightarrow$ control `Neu`<br>Treated `Prom1` $\not\rightarrow$ control `Alt` | Separation of condition-specific fates and loss of the `Alt` fate after ATRA. |
| Reachable fixed points | Control:<br>– From `Prom1`: `Rep`, `Neu`, `Alt`<br>– From `Prom2`: `Neu`, `Alt`<br>Treated:<br>– From `Prom1`: `Rep`, `Neu`<br>– From `Prom2`: `Neu` | Restriction of long-term behaviours reachable from promyelocytic macrostates. |
| | **Hematopoiesis** | |
| Fixed points and inequalities | Fixed points: S2, S4, S5<br>Inequalities:<br>– S2 $\neq$ S4<br>– S2 $\neq$ S5<br>– S4 $\neq$ S5 | Distinct lymphoid-primed, granulocyte-monocyte and megakaryocyte-erythroid endpoints. |
| Reachability and non-reachability | Reachability:<br>– S1 $\rightarrow$ S0 $\rightarrow$ S2<br>– S0 $\rightarrow$ S3 $\rightarrow$ S4<br>– S3 $\rightarrow$ S5<br>Non-reachability: S3 $\not\rightarrow$ S2 | Early differentiation and the lymphoid–myeloid and GMP–MEP bifurcations reconstructed by STREAM. |
| Reachable fixed points | – From S0: S2, S4, S5<br>– From S3: S4, S5 | Restriction of long-term behaviours to downstream lineage endpoints. |

**Supplementary Table 6:** Principal gene-selection and BN-inference settings and outputs. Certification refers to the configured optimisation objective. When important nodes are specified, the objective vector is lexicographic: the number of retained important nodes is maximised before the total number of retained nodes.

| Item | APL | Hematopoiesis |
|---|---|---|
| Prior regulatory domain | Mouse DoRothEA levels A–B | Mouse DoRothEA levels A–C |
| Initial domain size | 5,198 | 9,170 |
| Soft-stage retained domain | 5,016/5,198 (uncertified incumbent) | 9,113/9,170 (solver-certified objective vector) |
| Complete-constraint retained domain | 515/543 (feasible, uncertified incumbent) | 745/746 (feasible, uncertified incumbent) |
| Strong-constant scope | Soft constraints | Relaxed constraints |
| Strong constants removed | 4,473 | 8,367 |
| Post-constant domain | 543/5,016 (solver-certified within propagated domain) | 746/9,113 (solver-certified within propagated domain) |
| Final domain size | 515 | 745 |
| Maximum clause bound | 4 | 5 |
| Continuation | Clause and domain continuation during node selection | Clause and domain continuation at relaxed and seed stages |
| Random seed | 10 | 10 |
| Lock stage | Invoked with zero optimisation budget; no additional search | Invoked with zero optimisation budget; no additional search |
| BN inference strategy | Subset-minimal enumeration | Subset-minimal enumeration |
| Number of BNs retained | 1,000 | 1,000 |

**Supplementary Table 7:** Minimum, median and maximum numbers of minimal trap spaces and most-permissive reachable attractors per network across the 1,000 subset-minimal BNs retained for each case study. APL attractors were counted from the completed control and treated `Prom1` configurations and matched exactly to the corresponding terminal macrostates. Hematopoiesis attractors were counted from the completed `S1` and `S0` configurations.

| Quantity | Minimum | Median | Maximum |
|---|---|---|---|
| **APL** | | | |
| Minimal trap spaces | 6 | 6 | 6 |
| Attractors from control `Prom1` | 3 | 3 | 3 |
| matching control `Rep` | 1 | 1 | 1 |
| matching control `Alt` | 1 | 1 | 1 |
| matching control `Neu` | 1 | 1 | 1 |
| Attractors from treated `Prom1` | 2 | 2 | 2 |
| matching treated `Rep` | 1 | 1 | 1 |
| matching treated `Neu` | 1 | 1 | 1 |
| **Hematopoiesis** | | | |
| Minimal trap spaces | 3 | 3 | 3 |
| Attractors from `S1` | 3 | 3 | 3 |
| Attractors from `S0` | 3 | 3 | 3 |

**Supplementary Table 8:** Computational environment and wall-clock runtimes derived from the retained timestamped logs and, where indicated, configured timeout budgets. Durations are rounded to the nearest second.

| Item | APL | Hematopoiesis |
| --- | --- | --- |
| Operating system | Ubuntu 22.04.5 LTS; Linux 6.8.0-138-generic x86_64 | Same host |
| Processor | Intel Core Ultra 7 155H; 22 logical CPUs | Same host |
| Memory | 64 GB RAM; 50 GB cap | 64 GB RAM; 50 GB cap |
| Parallelism | Omics: 16 jobs<br>Node selection: up to 16 independent one-thread solves<br>BN enumeration: multi-threaded Clingo portfolio | Same configuration |
| Clingo | 5.8.0 | 5.8.0 |
| Logged upstream analysis | Matrix loading through consensus binarisation: 10 min 55 s<br>Potency scoring: 2 min 4 s | STREAM macrostate construction: 42 s<br>Binarisation: 38 s |
| Model specification | 5 s | 16 s |
| Gene-selection runtime | Soft: 15 h (timeout)<br>Strong constants: 52 s<br>Relaxed: 1 h 20 min 43 s<br>Seed: 5 h (timeout) | Soft: 5 min 48 s<br>Relaxed: 1 h 13 min 42 s<br>Strong constants: 1 h 13 min 5 s<br>Seed: 7 h 29 min 14 s (OOM kill) |
| BN-enumeration runtime | Approximately 2 h 34 min | 6 min 9 s |
| Cumulative reported runtime | Approximately 24 h 8 min | Approximately 10 h 10 min across retained-output stages; preprocessing before STREAM excluded |

**Supplementary Table 9:** Clingo solver configuration and source of parallelism for the historical comparison and the scBOLT procedure. One-thread node selection corresponds to the configuration historically selected by the authors after empirical benchmarks showed that it outperformed multi-threaded search in the large majority of tested satisfiable-node-maximisation instances.

| Procedure | Solver configuration | Parallelisation |
| --- | --- | --- |
| Historical node selection | One Clingo thread (`JOBS=1` override) | Single monolithic optimisation |
| scBOLT node selection | One Clingo thread per solve | Up to 16 independent domain-continuation subproblems |
| BN enumeration | Multi-threaded Clingo portfolio | Parallel enumeration |

**Supplementary Table 10:** Effects of constant perturbations of the 40 feedback-core genes across the 1,000 retained APL Boolean networks. Both constant values were evaluated for every gene. The table reports only the 32 perturbation directions, covering 31 distinct genes, with a non-zero score in at least one targeted phenotype-reachability outcome; all other directions are omitted. This filter does not use the off-target count. A value of 0 or 1 after a gene name denotes fixation to an off- or on-like transcriptomic state, respectively, rather than a direct protein activity measurement. Values are counts over BN–condition pairs. For each perturbation and pair, a targeted effect was counted only if all non-targeted baseline attractors remained reachable and no perturbed attractor was mapped to a previously inaccessible reference attractor. The off-target column counts BN–condition pairs in which a protected baseline attractor was lost or a perturbed attractor mapped to a previously inaccessible reference attractor. Neutral pairs with neither a targeted nor an off-target effect are not counted.

| | control | | treated | |
|---|---|---|---|---|
| perturbation | `Alt` preserved, `Rep` lost | `Alt` and `Rep` lost | `Rep` lost | off-target |
| Cebpe=1 | 1000 | 0 | 1000 | 0 |
| E2f1=0 | 1000 | 0 | 1000 | 0 |
| Egr1=0 | 1000 | 0 | 1000 | 0 |
| Esr1=0 | 1000 | 0 | 1000 | 0 |
| Foxa1=0 | 1000 | 0 | 1000 | 0 |
| Foxm1=0 | 1000 | 0 | 1000 | 0 |
| Myc=0 | 1000 | 0 | 1000 | 0 |
| Ppara=0 | 1000 | 0 | 1000 | 0 |
| Trp53=0 | 1000 | 0 | 1000 | 0 |
| Trp63=0 | 1000 | 0 | 1000 | 0 |
| Mycn=0 | 999 | 0 | 999 | 0 |
| Rela=1 | 973 | 0 | 1000 | 0 |
| Ets2=0 | 968 | 0 | 968 | 0 |
| Jun=0 | 958 | 0 | 958 | 0 |
| Stat3=1 | 1000 | 0 | 1 | 0 |
| Rara=1 | 0 | 0 | 1000 | 0 |
| Stat1=1 | 0 | 1000 | 0 | 0 |
| Stat2=1 | 0 | 1000 | 0 | 0 |
| Twist1=1 | 1000 | 0 | 0 | 0 |
| Nfkb2=1 | 0 | 0 | 986 | 0 |
| Yy1=0 | 22 | 0 | 22 | 0 |
| Gata3=1 | 7 | 0 | 8 | 0 |
| Pou2f2=1 | 10 | 0 | 5 | 0 |
| Spi1=1 | 10 | 0 | 5 | 0 |
| Tbp=0 | 4 | 0 | 4 | 0 |
| Vdr=0 | 4 | 0 | 4 | 0 |
| Ets1=1 | 7 | 0 | 0 | 0 |
| Mitf=1 | 7 | 0 | 0 | 0 |
| Gata3=0 | 6 | 0 | 0 | 739 |
| Gata1=1 | 0 | 0 | 2 | 0 |
| Runx1=1 | 0 | 0 | 2 | 0 |
| Esr2=1 | 0 | 0 | 1 | 0 |

**Supplementary Table 11:** Macrostate-specific binarisation rates in the hematopoiesis case study. Values are percentages of the 4,241 genes selected for binarisation that were assigned a Boolean value by scBoolSeq, differential-expression analysis (DEA), or their consensus.

| Macrostate | scBoolSeq (%) | DEA (%) | Consensus (%) |
|---|---|---|---|
| S0 | 4.5 | 13.2 | 14.8 |
| S1 | 6.4 | 21.9 | 23.6 |
| S2 | 7.8 | 15.2 | 20.4 |
| S3 | 7.4 | 5.2 | 11.5 |
| S4 | 14.0 | 34.3 | 39.3 |
| S5 | 9.2 | 15.6 | 21.5 |

**Supplementary Table 12:** Pairwise intersections between the normalised component sets of the Collombet [5], Hamey [6], Moignard [7], Chevalier [8] and scBOLT hematopoiesis models. Off-diagonal entries report the number of shared gene components, whereas diagonal entries report the total number of components in each model. The 39 components of the published Chevalier model induce a 137-interaction DoRothEA prior.

| | Pairwise component intersections | | | | |
|---|---|---|---|---|---|
| Model | Collombet | Hamey | Moignard | Chevalier | scBOLT |
| Collombet | **21** | 3 | 4 | 6 | 12 |
| Hamey | – | **31** | 13 | 6 | 16 |
| Moignard | – | – | **20** | 6 | 9 |
| Chevalier | – | – | – | **39** | 27 |
| scBOLT | – | – | – | – | **745** |

#### Supplementary Figures

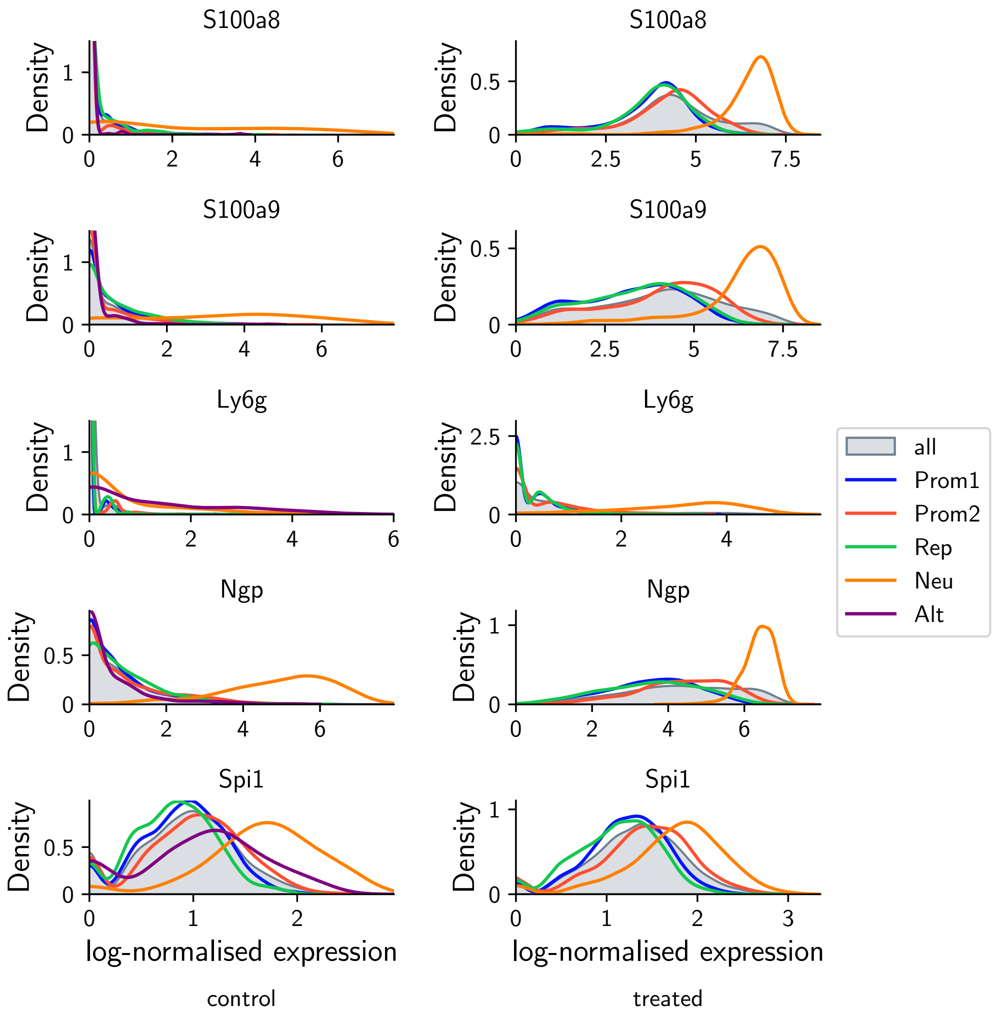


**Supplementary Figure 1:** Gene-expression density profiles for granulocytic markers in the APL dataset. Curves show log-normalised expression distributions for S100a8, S100a9, Ly6g, Ngp, and Spi1 across the annotated cell populations in the control and ATRA-treated conditions; grey distributions pool all retained cells within each condition. These profiles illustrate the distribution shapes used by scBoolSeq and the population-specific expression shifts used by differential-expression analysis during consensus macrostate binarisation.

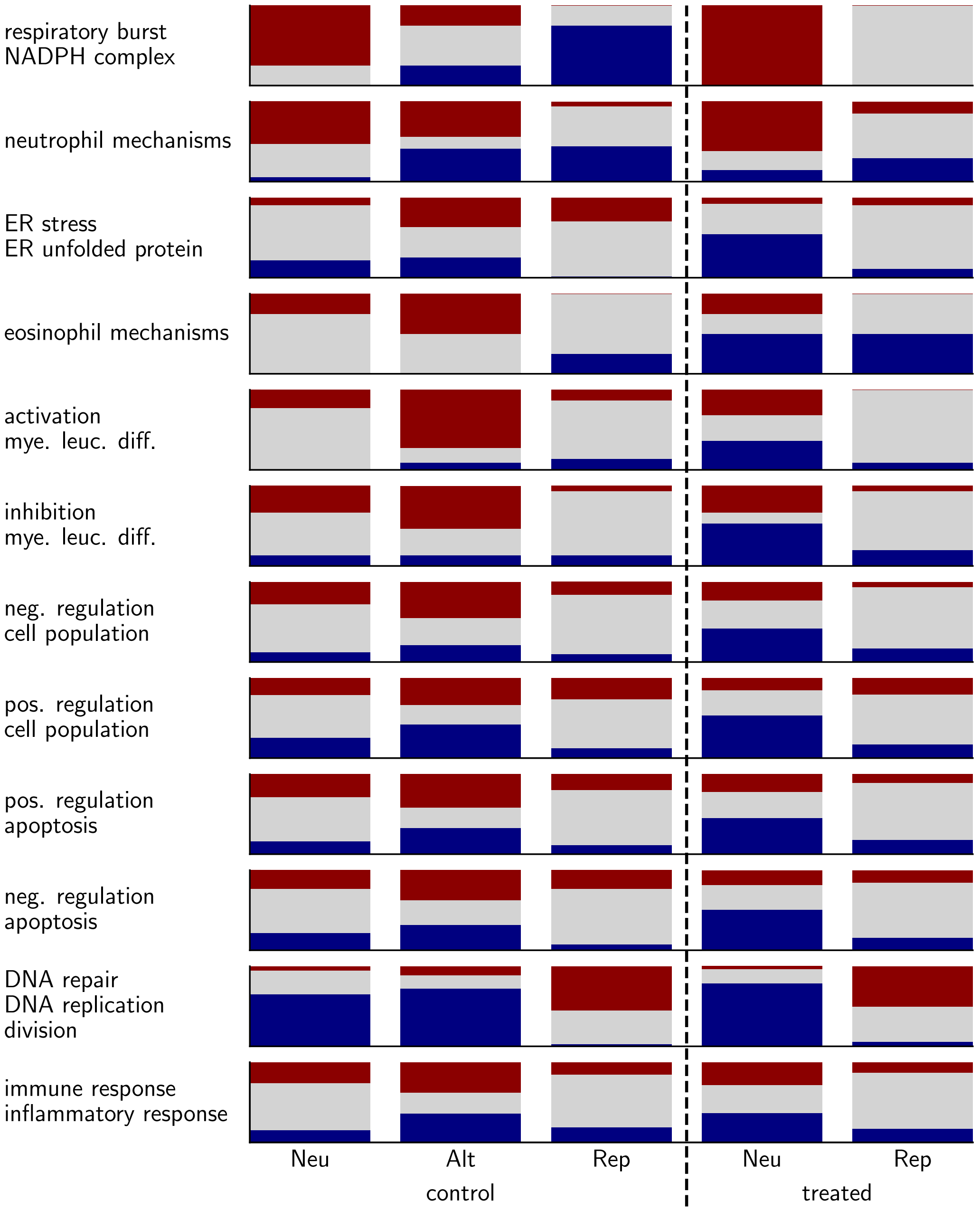


**Supplementary Figure 2:** Extended GO-derived functional profiles of binarised APL macrostates. For each functional gene set and terminal macrostate, stacked bars report the proportion of genes assigned to a low (0) transcriptomic state (navy), left unassigned (grey), or assigned to a high (1) transcriptomic state (dark red). The dashed line separates control and ATRA-treated macrostates. This exploratory panel extends the focused functional comparison presented in the main text.

**Supplementary Figure 3:** Ensemble-aggregated influence graph for the 1,000 retained APL BNs. Green and red edges denote activating and inhibitory influences, respectively; edge labels report the empirical fraction of the 1,000 retained BNs containing each signed influence, and line style also encodes this frequency. A frequency of one means that the influence is present in all 1,000 retained BNs, not that it is universal over the complete compatible solution space. To reduce the full 515-gene graph, non-feedback genes with identical signed predecessor and successor patterns in the same frequency bins were merged into composite nodes whose members are separated by vertical bars. Feedback genes were not collapsed, and isolated nodes were omitted. Node shading encodes the frequency of the most common DNF implicant structure across the ensemble, with stronger shading indicating greater structural stability.

**Supplementary Figure 4:** Ensemble-aggregated influence graph for the 1,000 retained BNs in the hematopoiesis case study. Green and red edges denote activating and inhibitory influences, respectively; edge labels report the empirical fraction of the 1,000 retained BNs containing each signed influence, and line style also encodes this frequency. A frequency of one means that the influence is present in all 1,000 retained BNs, not that it is universal over the complete compatible solution space. To reduce the full 745-gene graph, non-feedback genes with identical signed predecessor and successor patterns in the same frequency bins were merged into composite nodes whose members are separated by vertical bars. Feedback genes were not collapsed, and isolated nodes were omitted. Node shading encodes the frequency of the most common DNF implicant structure across the ensemble, with stronger shading indicating greater structural stability.

### Supplementary Data and Code

- **APL workflow.** The complete executable configuration and dynamical specification are provided in `apl/params-gsm.yml` and `apl/spec.yml`, respectively. Generated macrostates, partial Boolean states, retained regulatory domains and inferred BNs are stored under `apl/project_gsm/` or can be regenerated from these entry points. Annotation-support result tables are stored under `apl/results/annotation/`.
- **Hematopoiesis workflow.** The corresponding files are `hematopoiesis/config/params-abc.yml` and `hematopoiesis/config/spec.yml`; generated outputs are stored under `hematopoiesis/project/`.
- **Figure-generation notebooks.** Transcriptomic analyses and Boolean-network analyses are documented in the `notebooks/omics.ipynb` and `notebooks/bn.ipynb` files within each case-study directory.
- **Versioned archive.** All configurations, scripts and notebooks are available from https://github.com/bnediction/scbolt-case-studies and archived at https://doi.org/10.5281/zenodo.22959149.
