## Supplementary figures and images for "Scalable and semi-automated Boolean network inference from multi-condition single-cell transcriptomes with scBOLT"

### Supplementary Figure 3

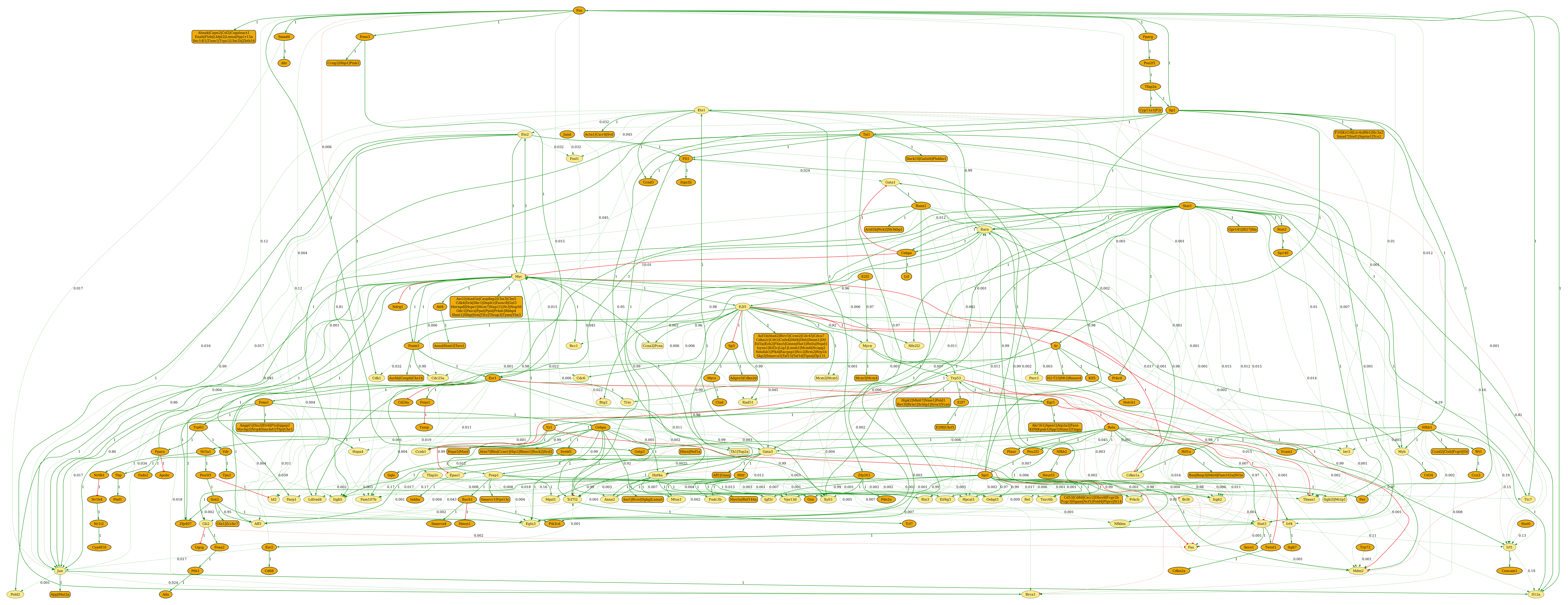

### Supplementary Figure 4

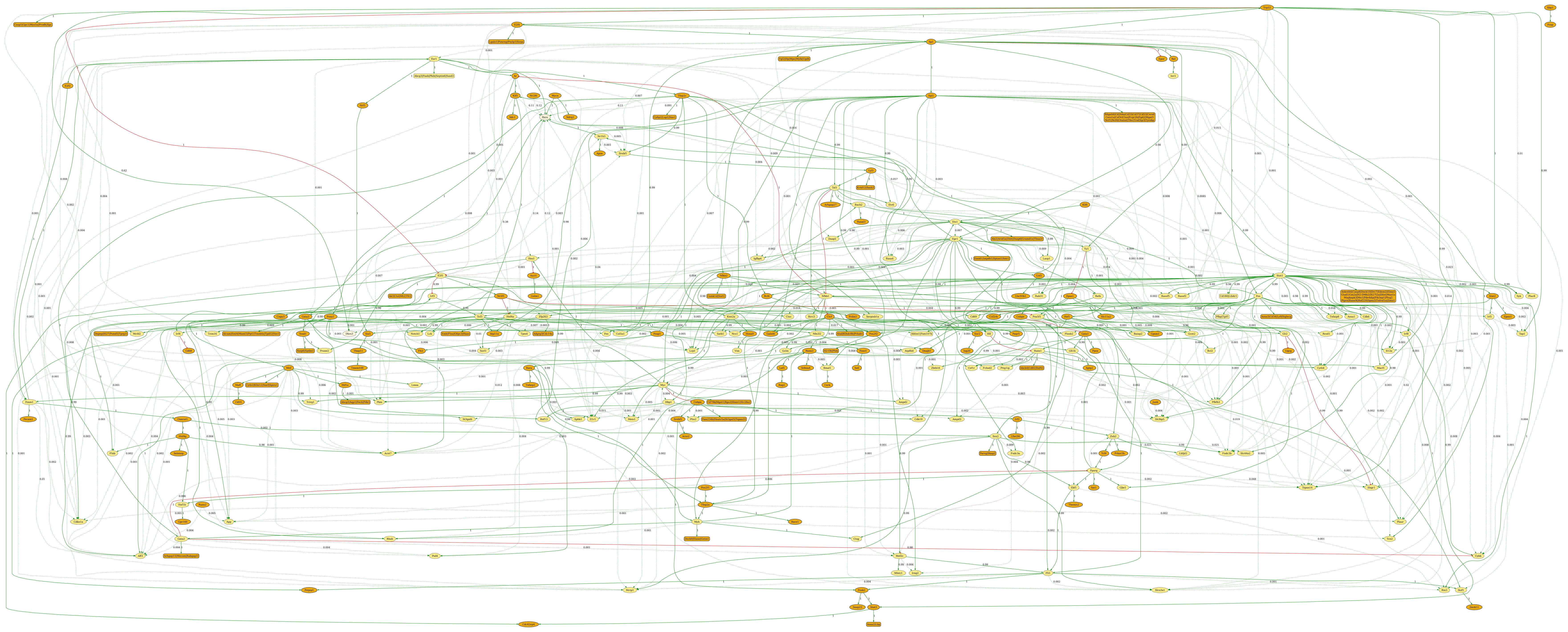
